# Automated pup-level analysis reveals distinct effects of prenatal CBD and THC exposure on maternal retrieval

**DOI:** 10.64898/2026.09.22.753513

**Authors:** Alba Cáceres-Rodríguez, Benjamin Strauss, Julien Ferragu, Jessica Pereira, Daniela Iezzi, Pascale Chavis, Olivier J. Manzoni

**Affiliations:** INMED, INSERM, Aix-Marseille University, Marseille, France

**Keywords:** in utero, cannabidiol, delta9-tetrahydrocannabinol, cannabis, pup retrieval, maternal behavior, deep learning

## Abstract

This study examined how prenatal CBD and Δ9-tetrahydrocannabinol (THC) exposure affects maternal caregiving in C57BL/6J mice. We developed the Machine-Automated Scoring of the Pup Retrieval Test (MAS-PRT) to overcome limitations of manual behavioral scoring. MAS-PRT integrates Multi-Animal DeepLabCut for dam and pup tracking with Detectron2 for dynamic nest reconstruction. Validated against 170 manually annotated retrieval trials, the pipeline showed high concordance with manual measurements and enabled reproducible extraction of encounter latency, retrieval latency, and locomotor trajectories.

Prenatal exposure did not impair general nest-building or overall home-cage maternal care. However, pups from both CBD and THC groups showed reduced body weight at postnatal day 5. Cox proportional hazards modeling revealed divergent effects by compound: CBD-exposed dams exhibited a weight-dependent increase in probability of encountering and retrieving lighter pups, independent of pup sex. Spatial tracking further showed that dams traversed significantly shorter total trajectories when retrieving female progeny exposed to either compound. Longitudinal trial-by-trial analysis indicated intact task acquisition in controls and CBD dams, whereas THC dams displayed a flattened learning curve driven by lower retrieval latencies on initial trials. Together, these findings indicate that prenatal cannabinoid exposure does not produce generalized disruption of maternal care but instead induces compound-specific alterations in maternal reactivity, retrieval kinematics, and learning dynamics.

## Introduction

Prenatal exposure to cannabis poses significant risks to both maternal well-being and fetal development (Koto et al., 2022; Navarrete et al., 2020; Scheyer et al., 2019). As the most commonly used illicit substance among pregnant and breastfeeding women (Metz et al., 2017; Scheyer et al., 2019; Young-Wolff et al., 2019), cannabis compounds, specifically cannabidiol (CBD) and delta-9-tetrahydrocannabinol (THC), readily cross the placental barrier due to their lipophilic nature, accumulating in fetal tissues (Maciel et al., 2022; Sarrafpour et al., 2020). While THC is the primary psychoactive component, recent years have seen a surge in the use of CBD, a non-intoxicating derivative of *Cannabis sativa*, by expectant mothers seeking relief from pregnancy-related discomforts such as anxiety, nausea, and insomnia (Bhatia et al., 2024; Sarrafpour et al., 2020; Volkow et al., 2019). Despite this growing trend, systematic research regarding the specific effects of CBD on maternal care remains scarce (Compagno et al., 2025; DeVuono et al., 2024; Swenson et al., 2023).

Maternal care is a critical determinant of offspring development and is shaped by the coordinated engagement of sensory, motivational, and motor processes (Bridges, 2015; Curley & Champagne, 2016). In humans, substance use during pregnancy is frequently associated with atypical parenting behaviors that disrupt mother-infant attachment (Cataldo et al., 2019; Parolin & Simonelli, 2016). Neurobiologically, the transition to motherhood involves profound remodeling of regulatory circuits, emotional processing, and reward systems (Curley & Champagne, 2016); circuitry that can be also modified by drug exposure (Gammie, 2005), suggesting that prenatal cannabinoid exposure could interfere with the neural mechanisms governing parental behavior Poor maternal care in rodents has been directly linked to neurodevelopmental deficits and neuropsychiatric vulnerabilities in offspring (Bridges, 2015), highlighting the necessity of examining parent-infant interactions in the context of drug exposure.

Existing studies of prenatal cannabinoid exposure have primarily relied on conventional behavioral measures and manual scoring. Studies using low-to-moderate prenatal THC exposure (2-5 mg/kg) have generally reported preserved maternal behavior, including nest building and pup-directed behaviors (Bara et al., 2018; Frau et al., 2019; Traccis et al., 2020). However, emerging evidence suggests that while prenatal CBD exposure may not immediately abolish maternal behaviors, it can influence offspring outcomes, such as protecting against OCD-like behaviors when offspring are cross-fostered to drug-free mothers (Compagno et al., 2025).

In rodents, pup retrieval test (PRT) provides a particularly useful assay of maternal responsiveness because it requires the dam to detect a displaced pup, approach it, and return it to the nest. Importantly, maternal retrieval is influenced by properties of the pup, including body weight and sex of the pup and ultrasonic vocalizations, as well as by the dam’s previous experience with the task (Bowers et al., 2013; Ehret, 2005; Weber & Olsson, 2008; Winters et al., 2023). However, these studies often rely on manual scoring methods that may lack the sensitivity to detect subtle, trial-level variations in maternal reactivity or fail to account for confounding variables like pup weight and sex.

Here, we investigated the effects of prenatal CBD and THC exposure on maternal behavior using complementary measures of nest building, home-cage nursing, and pup retrieval. We further developed the Machine-Automated Scoring of the Pup Retrieval Test (MAS-PRT), an automated pipeline integrating multi-animal DeepLabCut tracking with dynamic nest detection to quantify individual pup encounters, retrievals, and maternal trajectories. We reasoned that automated, pup-level analysis combined with time-to-event modeling could resolve behavioral features that may be obscured by aggregate measures of mean dam retrieval performance. We therefore examined maternal responses as a function of pup treatment, body weight, sex, trial order, and retrieval trajectory. This approach revealed that prenatal CBD and THC exposure produced distinct alterations in maternal retrieval dynamics despite largely preserved conventional measures of maternal care.

## MATERIALS AND METHODS

### Animals and Housing

All experimental procedures adhered to the European Communities Council Directive (86/609/EEC) and the United States NIH Guide for the Care and Use of Laboratory Animals. The project was authorized by the French Ethical Committee (APAFIS#49376). Adult male and female C57BL/6J mice (6–8 weeks old) were purchased from Charles River. Animals were housed in standard lid-topped Plexiglas cages (42 × 27 × 14 cm) under controlled conditions: temperature 21 ± 1°C, relative humidity 60 ± 10%, and a 12-hour light/dark cycle (lights on at 07:00). Food and water were available *ad libitum*.

Following a one-week acclimation period, females were paired with a single male in the late afternoon. The presence of a vaginal plug was designated as gestational day 0 (GD0). Pregnant dams were then individually housed. Postnatal day 0 (PND0) was defined as the day of birth. Litters were not culled at birth to preserve natural litter size variability. Pups were weaned on PND21 and subsequently housed separately by sex.

### Prenatal Drug Exposure

From GD5 to GD18, pregnant dams received daily subcutaneous (s.c.) injections of either a vehicle solution, 3 mg/kg of CBD, or 3 mg/kg of THC(Iezzi et al., 2022, 2025; Maciel et al., 2022). The cannabinoids were obtained from the NIDA Drug Supply Program. Both compounds were dissolved separately in a vehicle solution consisting of Cremophor-EL (Sigma-Aldrich), ethanol, and saline in a 1:1:18 ratio, administered at a volume of 4 mL/kg. Control dams (“Sham”) received an equivalent volume of the vehicle solution alone.

### Behavioural assesments

#### Nest Building Skills

Nest quality was assessed daily at 09:30 am. Dams were provided with 11 g of pressed aspen wood wool from the onset of gestation. Nests were manually scored using a modified version of Deacon’s 5-point scale(Deacon, 2006; Hess et al., 2008), ranging from 0 to 4 (Kuroda & Tsuneoka, 2013): 0, no shredding and no visible nest site; 1, material shredded completely or partially with an identifiable but flat nest; 2, material shredded and a saucer-shaped nest formed; 3, material completely shredded with walls tall enough to cover the animal; and 4, material completely shredded forming a fully enclosed nest with a complete roof covering the animal.

#### Home-Cage Dam Behavior

Maternal behavior was observed daily from PND1 to PND10 at 10:00 for 20 minutes, right after the nest assessment. As most of the nests were partially or fully enclosed, prior to the test, the top of the nest was manually enlarged to provide better visibility for the experimenter. After opening the cage to open the nest, the dams were allowed to habituate for 15 minutes before testing with the transparent lid closed. During each session, the dam and litter were observed for five 60-second intervals, resulting in a total observation time of 300 seconds per day per litter.

Observations focused on nest occupancy, which recorded whether the dam was inside or outside the nest, with time spent outside (eating, drinking, and grooming) noted but not quantified. Additionally, nursing behavior was measured as the time spent inside the nest and classified into two categories: active nursing, defined as the dam arched over pups with rigid limbs and head depressed, and passive nursing, defined as the dam lying flat on the litter with minimal limb support.

#### Pup Retrieval Test

Maternal reactivity was assessed on PND5, following a protocol adapted from(Winters et al., 2022). Testing occurred between 10:30 and 13:00 in a custom soundproofed plastic arena equipped with a top-mounted camera (Foscam C2 IP camera; 2046 × 2046 resolution, 15 fps) positioned 50 cm above the floor.

For the testing procedure, the home cage was moved to the testing room, and the dam was allowed to acclimate for 45 minutes with the nest undisturbed and the lid of the cage open. Subsequently, one pup was removed, weighed, sexed (Wolterink-Donselaar et al., 2009), marked, and placed in a heated glass beaker (35°C). Once the dam was confirmed to be in the nest, the isolated pup was placed in the corner of the cage furthest from the nest. The dam was given 90 seconds to retrieve the pup; if retrieval did not occur, the pup was returned to the nest and the next littermate was tested, with trials failing to occur within 90 seconds classified as censored events (failure). Recorded metrics included the time to first encounter (tFE), defined as the time from the start until the dam contacted the pup; the time to retrieve (tR), defined as the total time from the start until the pup was returned to the nest; and success, recorded as a binary outcome where 1 indicated retrieval within 90 seconds and 0 indicated no retrieval.

#### Automated Analysis Pipeline (MAS-PRT)

Videos were processed using a custom deep learning pipeline, the Machine-Automated Scoring of the Pup Retrieval Test (MAS-PRT). The workflow involved three stages: (1) tracking dam and pup movements, (2) automatic nest detection, and (3) data extraction (Suplementary Figure 1). Videos were pre-processed using the open-source video transcoder HandBrake (cropped to the home cage, resized to 720p, and converted to grayscale).

The model and instructions for adapting it to specific experimental setups are available online on GitHub: https://mas.readthedocs.io/en/latest/index.html

For the (1) *<u>tracking and pose estimation</u>*, we utilized Multi-Animal DeepLabCut (DLC) (Lauer et al., 2022). Seven key points were labeled for the dam and five for each pup (Figure 5A–B). The model was trained on 988 frames (selected via k-means clustering) using a ResNet50 backbone. Training continued for 40,000 iterations until performance plateaued. The dataset was split 95% for training and 5% for validation. 30 DLC tracked videos were randomly selected for manual verification of the tracking to ensure the was no swift in the identity of pup and dam. (2) *<u>Nest detection</u>* was performed using Detectron2 (Wu et al., 2019). The model was trained on 66 annotated frames from 33 videos. During analysis, 20 pseudo-randomly selected frames per video were used to generate a polygonal representation of the nest. An average polygon was computed per video to account for dynamic changes in nest shape during the test. Validation tests confirmed that 20 frames provided accuracy comparable to a 100-frame benchmark (Supplementary Figure 2). For each experiment analyzed, we manually reviewed the nest segmentation in the 20 randomly selected frames for nest detection in each video. (3) *<u>Data extraction and validation:</u>* The pipeline integrated DLC and Detectron2 outputs to extract metrics including: PRT success, tR, tFE, distance to first encounter (dFE), distance from first encounter to retrieval (dFE-R), total distance moved (dT), and nest area (cm²). The system was validated against manual annotations from 170 trials (from 170 different pups, all treatments represented), demonstrating high concordance with minimal deviation in latency metrics. The manual annotation was done one time by an experience researcher blind to the treatment. (Figure 5D).

### Statistical Analysis

All statistical analyses were performed using GraphPad Prism (v11) and Python (3.12). Gestational parameters were evaluated using the Kruskal-Wallis test and pup’s body weight two-way ANOVA followed by Tukey’s post hoc multiple comparison test. Nest quality and home-cage behavioral data were analyzed using linear mixed-effects models with Geisser-Greenhouse correction, designating treatment and time as fixed effects and subject as a random effect to account for repeated measures; Tukey’s post-hoc test was applied for multiple comparisons where appropriate. For the pup retrieval test (PRT), given the right-censored nature of the data (90-second upper limit), time-to-event (survival) analyses were conducted using a Cox proportional hazards regression (Cox-PH) model to predict first encounter and retrieval latencies. Covariates included pup sex, trial order, treatment, pup weight, and the interaction term (pup weight × treatment), with dam clustering and robust errors correction incorporated to account for within-litter correlations (litter effects). Equality of variances in pup weights across trial orders was verified using Levene’s test implemented in Python via the SciPy library (Supplementary Table 1). Longitudinal learning curves and trial-by-trial task acquisition were evaluated using LMMs modeling latency as a function of trial order and treatment, specifying SHAM as the reference category. Random effects included both a random intercept and a random slope for Trial grouped by Litter ID to account for individual litter trajectories over time. All Python modeling, statistical testing, and visualization were performed using statsmodels (LMMs and Benjamini-Hochberg/Bonferroni FDR adjustments), SciPy (scipy.stats for linear regressions, Levene’s test, and Kruskal-Wallis tests), pandas, NumPy, Seaborn, and Matplotlib. All statistical tests were two-tailed, and significance was set at *p* < 0.05.

## RESULTS

### Prenatal cannabinoid exposure does not modify gestational outcomes

The endocannabinoid (eCB) system plays a critical role in reproductive physiology, regulating processes such as fertilization, implantation, and neurodevelopment (Harkany et al., 2007; Innocenzi et al., 2019). Consequently, exposure to cannabis, its primary phytocannabinoids (THC, CBD), or synthetic agonists has been linked to adverse reproductive outcomes, including impaired fertility (Innocenzi et al., 2019), reduced ovarian reserve (Castel et al., 2020), and placental dysfunction leading to restricted fetal growth (El Marroun et al., 2009; Innocenzi et al., 2019; Rokeby et al., 2023). Disruption of eCB signaling can further compromise key events such as embryo transport (Wang et al., 2004) and uterine implantation (Schmid et al., 1997). Notably, high-dose oral CBD administration during gestation has previously been associated with reduced pup survival from birth to weaning (Compagno et al., 2025).

Prenatal exposure to CBD or THC did not significantly alter maternal gestational weight gain, litter size, or sex ratio at birth (Figure 1). All litters survived to weaning, and no overt maternal or reproductive toxicity was observed under the exposure conditions used here. These findings indicate that prenatal exposure to 3 mg/kg CBD or THC from GD5 to GD18 did not substantially alter the major gestational outcomes measured in this study.

**Figure 1:**
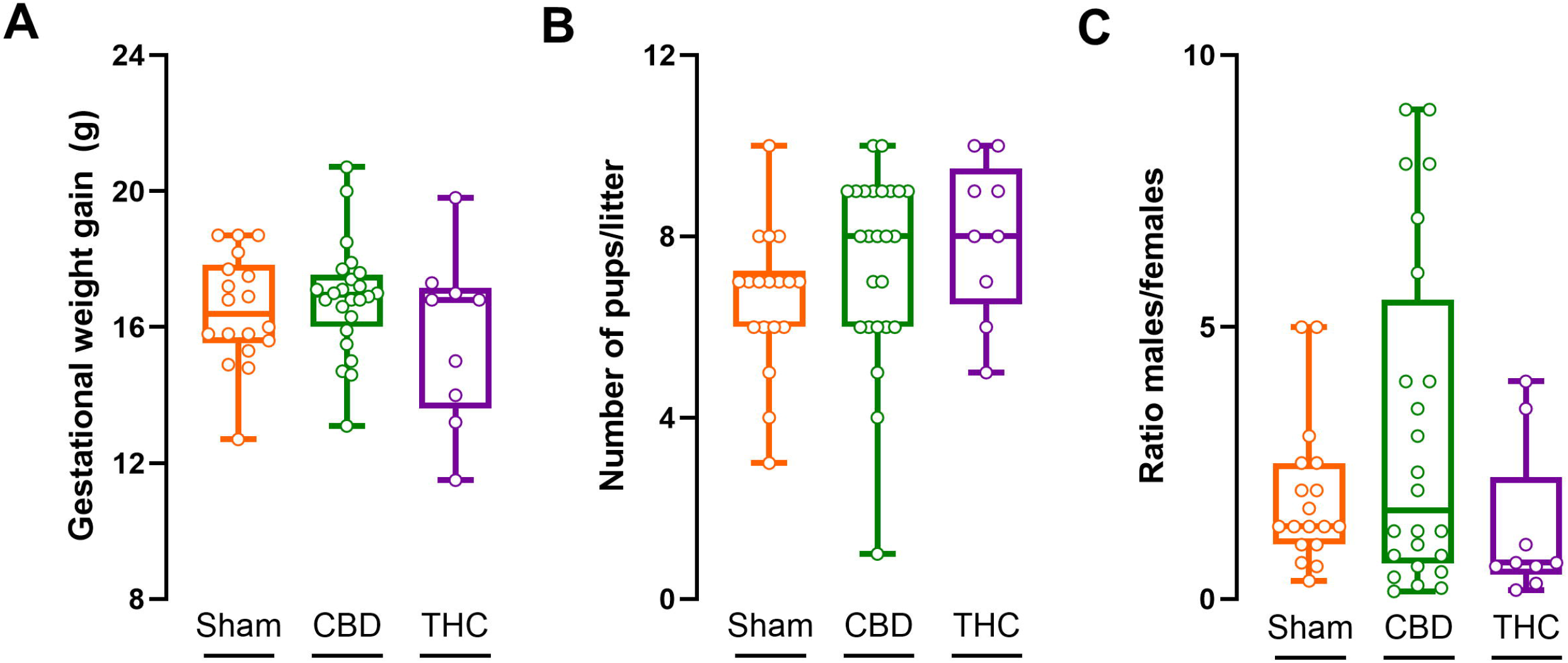
Prenatal CBD and THC exposure does not alter maternal weight gain or reproductive outcomes. (A) Maternal weight gain throughout gestation. (B) Litter size at birth. (C) Offspring sex ratio. Prenatal exposure to CBD or THC had no significant effect on maternal physiological or reproductive parameters. Data are presented as box-and-whisker plots showing median, minimum, and maximum values; each point represents a single litter (N= Sham 18 litters; CBD 25 litters; THC 8 litters). Statistical comparisons were performed using the Kruskal-Wallis test: H _(weight gain, 2)_= 1.601, p-value = 0.449; H_(#pups, 2)_= 5.02, p-value = 0.081; H_(ratioMF, 2)_= 3.548; p-value = 0.169. Only results with *p* < 0.05 were considered statistically significant and reported.

### Prenatal CBD and THC exposure reduces body weight at postnatal day 5 in pups independent of sex

Cannabis use during pregnancy has been associated with adverse outcomes, including fetal growth restriction and low birth weight (El Marroun et al., 2009; Oke et al., 2021). Specifically, perinatal exposure to CBD has been shown to influence the metabolic health of offspring (Compagno et al., 2025; Vanin et al., 2023). Pups prenatally exposed to either CBD or THC exhibited significantly lower body weight at PND5 compared with control pups (Figure 2). This growth deficit was observed consistently in both male and female pups, indicating that the impact of prenatal cannabinoid exposure on early growth is shared in both sexes of the progeny. Body-weight differences were no longer apparent by adulthood (Table 1), indicating that the early postnatal growth phenotype was transient under the conditions examined here.

**Figure 2:**
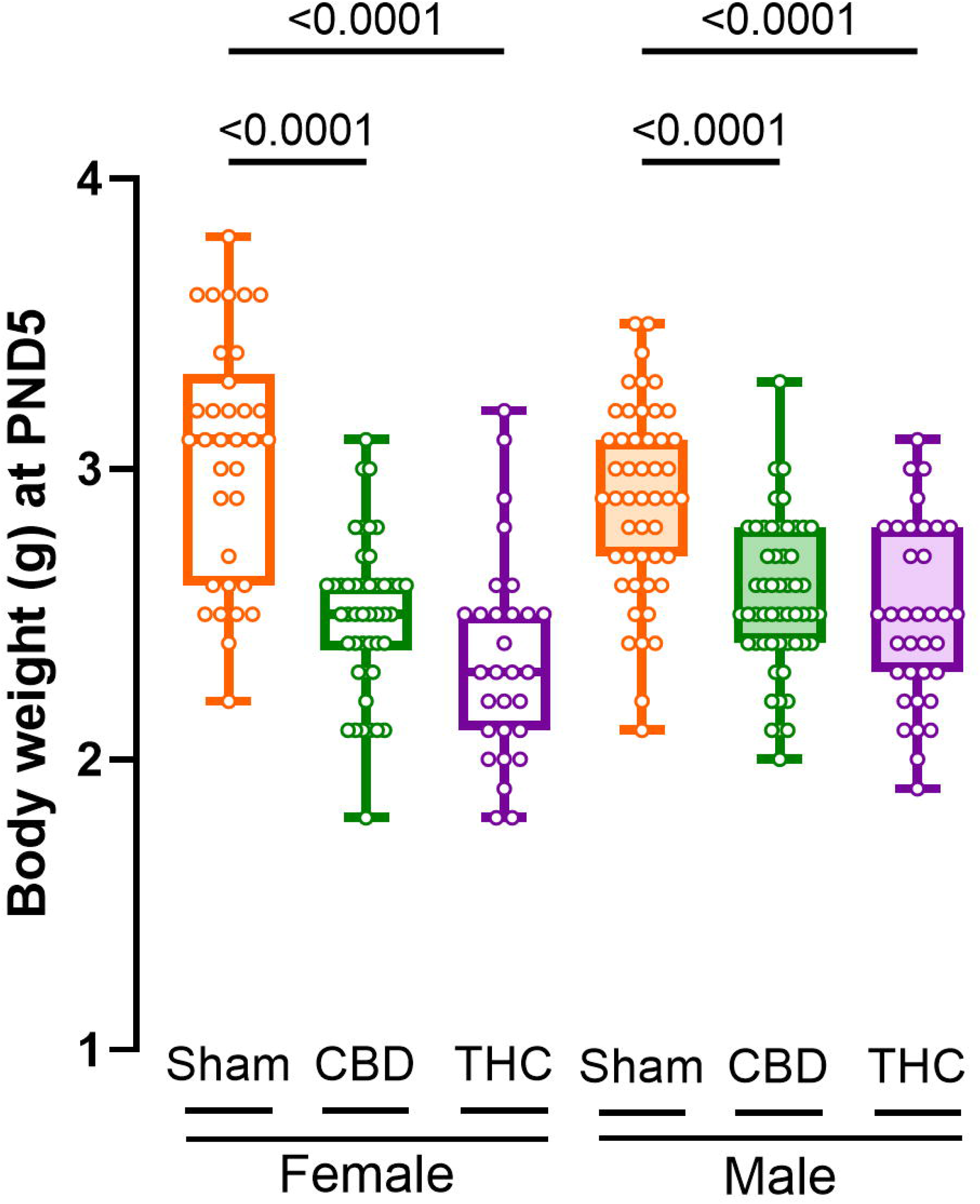
Prenatal CBD and THC exposure causes a sex-independent reduction in body weight at PND 5. Analysis of body weight by sex reveals that the growth restriction is consistent across males and females from CBD- and THC-exposed groups. Data are presented as mean ± SEM. Group sizes: Sham (34 females, 48 males), CBD (46 females, 54 males), and THC (29 females, 35 males). Statistical analysis: two-way ANOVA followed by Tukey’s post hoc multiple comparison test (F _(interaction 2, 240)_ = 3.801, p-value = 0.0237), F _(sex 1, 240)_ = 0.294, p-value = 0.588; F _(treatment 2, 240)_ = 65.98, p-value <0.0001). Only results with *p* < 0.05 were considered statistically significant and reported.

**Table 1:**
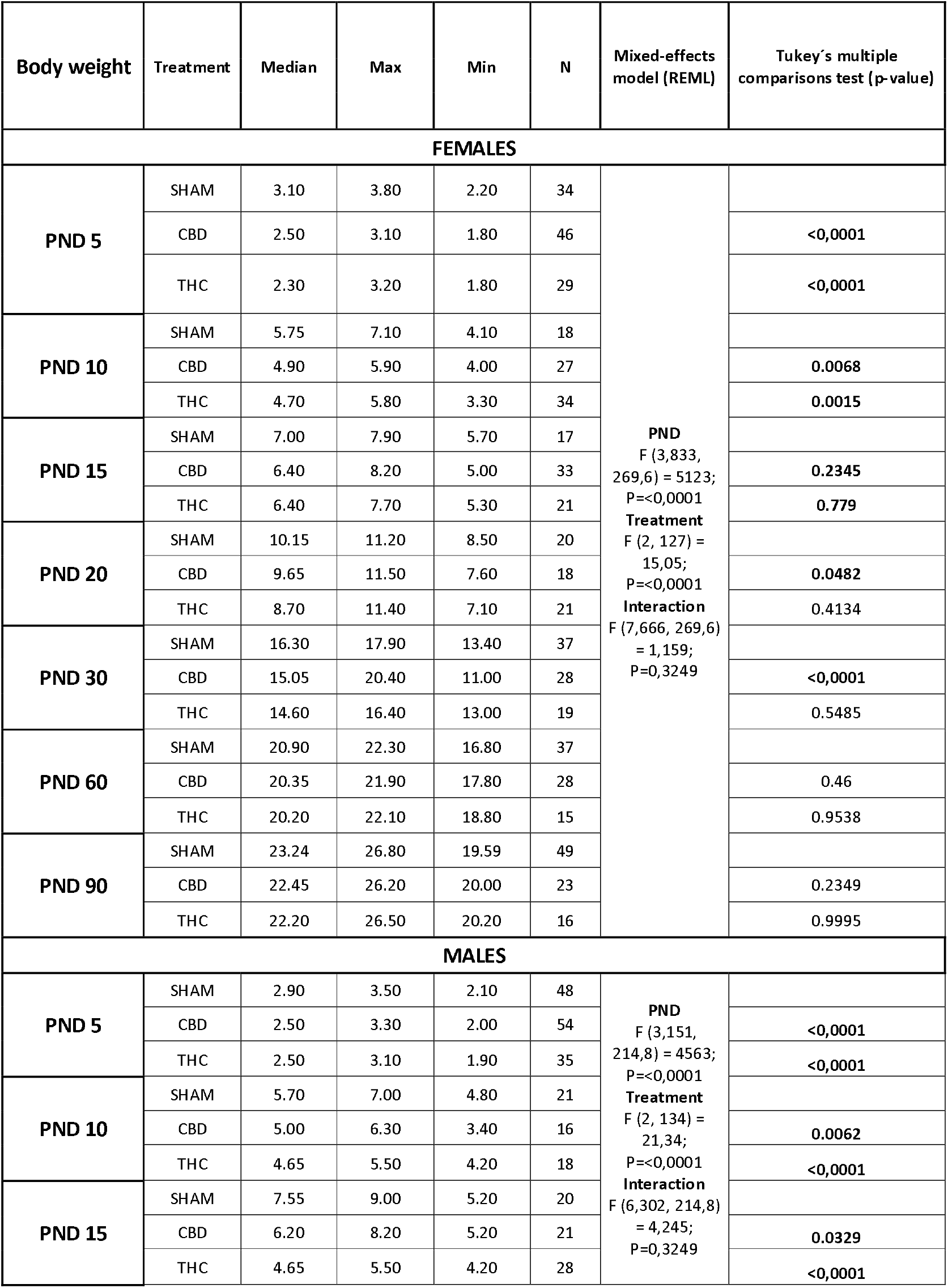
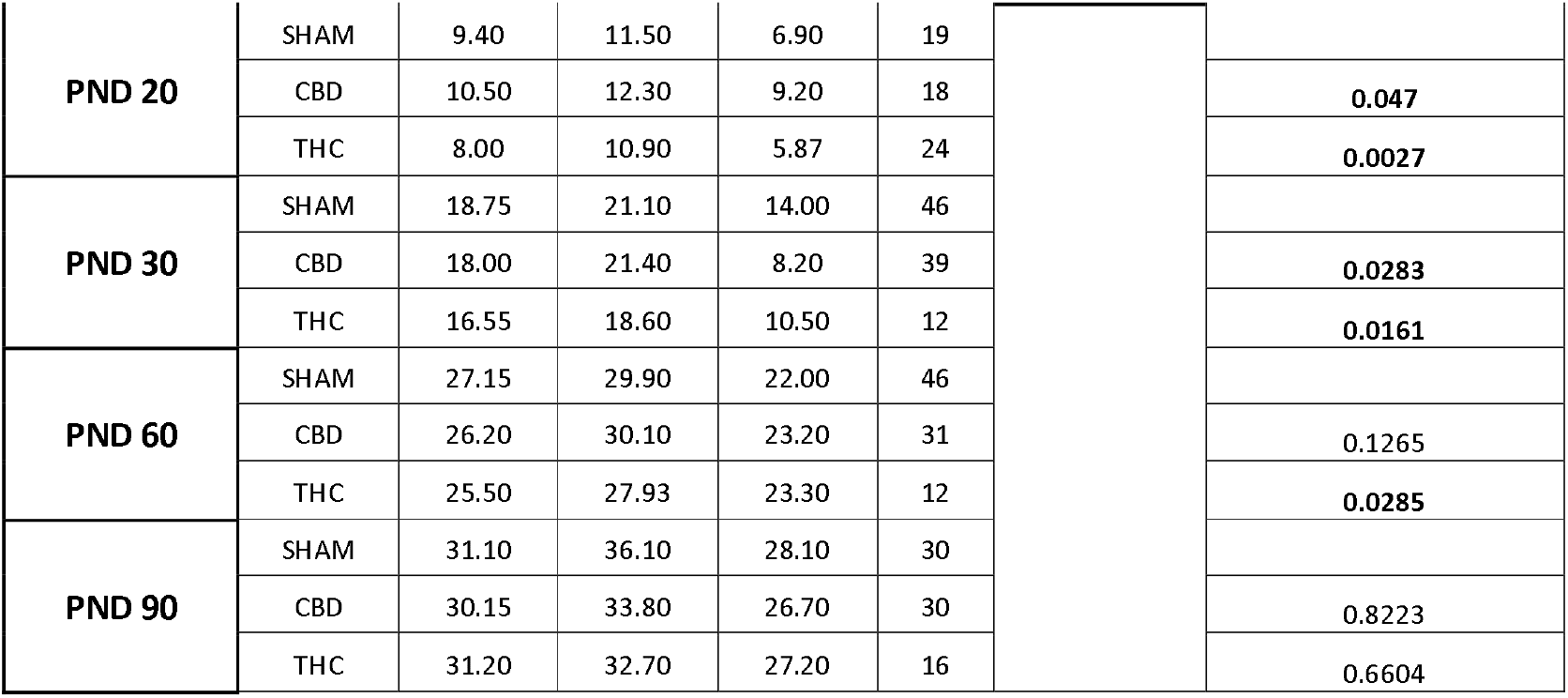
Body weight follow-up from early postnatal period till adulthood. Summary of body weight statistics (median, max, min, n) for females and males across treatments (Sham, CBD, THC) from PND 5 to PND 90. Statistical comparisons via Mixed-effects model (REML) for longitudinal trends and Tukey’s multiple comparisons test for pairwise treatment differences at each time point. *P*-values < 0.05 indicate significant differences from Sham controls.

### Nest-building behavior remains intact following prenatal cannabinoid exposure

In the wild, pregnant females construct complex nests to ensure offspring survival by providing essential protection and thermal insulation (Lisk et al., 1969; Tagawa et al., 2025; Topilko et al., 2022).

To determine whether prenatal cannabinoid exposure altered basic maternal behaviors, we first assessed nest construction and daily home-cage interactions with the litter.. Dams in all treatment groups successfully constructed nests of high complexity, consistently achieving scores of 3-4. Throughout the first 10 days of the postnatal period, these dams built fully enclosed nests that adequately covered their litters, ensuring necessary protection (Figure 3A; Supplementary Figure 3).

**Figure 3:**
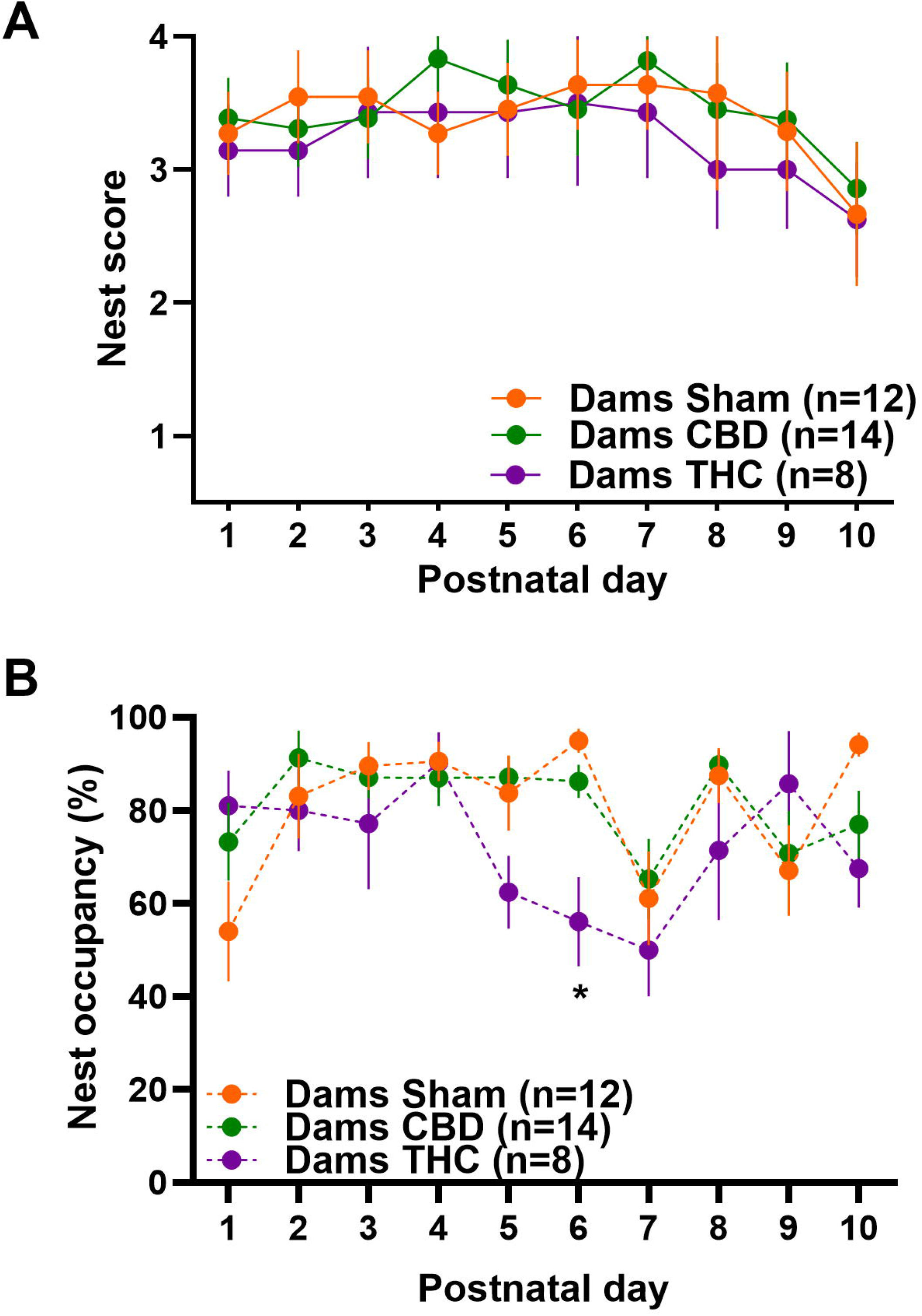
Prenatal THC, but not CBD, exposure transiently alters maternal nest occupancy. (A) Daily assessments of nest quality indicate that prenatal CBD or THC exposure does not disrupt nest-building behavior, which remains comparable to vehicle-treated controls. (B) Daily 20-min home-cage observations reveal a consistent preference for nest occupancy during the early postnatal period; however, THC-exposed dams exhibit a transient reduction in nest occupancy at PND 6. Data are expressed as mean ± standard error of the mean (SEM). Statistical analysis was performed via mixed-effects models with the Geisser-Greenhouse correction, followed by Tukey’s post hoc multiple comparison test: A (F _(time, 5.26, 127.9)_ = 6.417, p-value<0.0001; F _(treatment, 2, 29)_ = 2.370, p-value = 0.113, F _(interaction 10.52, 127.9)_ = 1.066, p-value = 0.393), B (F _(PND 5.08, 152.4)_ = 3.743, p-value = 0.003; F _(treatment 2, 31)_ = 2.107, p-value = 0.139; F _(interaction 10.16, 152.4)_ = 1.761, p-value = 0.03). Only results with *p* < 0.05 were considered statistically significant and reported.

Concurrent with nest assessments, daily home-cage observations (20 minutes/day) revealed that dams across all groups predominantly remained in the nest during daylight hours. A single exception was observed in the THC-exposed group on postnatal day 6, which exhibited a transient reduction in nest occupancy compared to controls (Figure 3B). Thus, prenatal CBD or THC exposure did not measurably disrupt nest-building behavior and nest occupancy.

### Prenatal cannabinoid exposure preserves high nest occupancy and normal nursing patterns

Pup-directed behaviors, such as nursing and licking, occur primarily within the nest, where the dam’s presence is essential for pup protection and healthy development (Bailey & Isogai, 2022; Lisk et al., 1969; Nowak et al., 2000). Nursing is particularly critical for offspring survival, providing both nutrition and thermoregulation(Nowak et al., 2000; Schuster et al., 2022).

In our study, dams across all treatment groups spent more than half of their time in the nest (Figure 3B), with most of this time dedicated to nursing (Figure 3A). Over the first 10 postnatal days, females consistently displayed a strong preference for nursing behaviors (Figure 4A). While generally stable, we observed a transient reduction in nursing among THC-exposed dams on postnatal day 6, which normalized by day 7.

**Figure 4:**
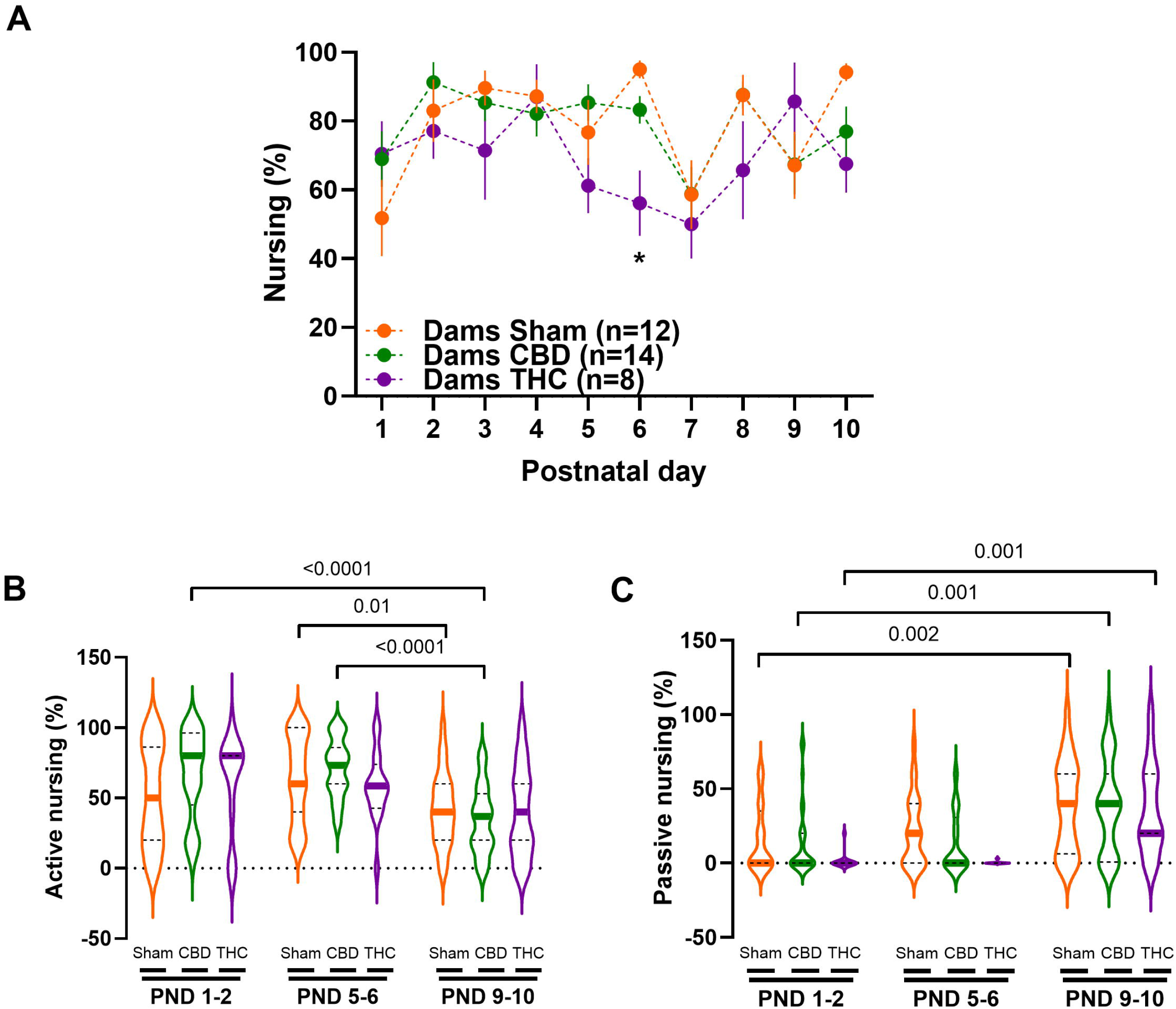
Prenatal THC exposure transiently alters nursing duration, while nursing style shifts from active to passive over postnatal development. (A) Time-course analysis of daily nursing duration across PND 1–10. THC-exposed dams exhibit a transient reduction in nursing duration at PND 6, concurrent with the nest occupancy deficit (see Figure 2B); otherwise, nursing duration remains consistent across groups. (B-C) Nursing style classification, showing the predominance of (B) active nursing in the early postpartum period and the transition to (C) passive nursing by the end of the follow-up (PND 9–10). (A) Data are presented as mean ± SEM in XY plots or as (B–C) violin plots indicating minimum, maximum, and median values. Statistical analysis was performed via mixed-effects models with the Geisser-Greenhouse correction, followed by Tukey’s post hoc multiple comparison test: A (F _(PND 5.121, 153.6)_ = 4.151, p-value = 0.001; F _(treatment 2, 31)_ = 2.293, p-value = 0.118; F _(interaction 18, 270)_ = 1.613, p-value = 0.056). B (F _(PND 2, 127)_ = 16.08, p-value <0.0001; F _(treatment 2, 65)_ = 0.4829, p-value = 0.619; F _(interaction 4, 127)_ = 1.496, p-value = 0.207). C (F _(PND 1.757, 111.6)_ = 35.25, p-value <0.0001; F _(treatment 2, 65)_ = 2.654, p-value = 0.078; F _(interaction 4, 127)_ = 1.143, p-value = 0.339). Only results with *p* < 0.05 were considered statistically significant and reported.

Furthermore, the profile of nursing behaviors shifted naturally over time. During the first postnatal week, dams primarily engaged in active nursing (Figure 4B; Supplementary Figure 4A). By postnatal day 10, passive nursing had become the dominant behavior (Figure 4C; Supplementary Figure 4B). Overall, these observations indicate that prenatal CBD or THC exposure did not produce a generalized disruption of maternal behavior under standard home-cage conditions.

### Machine-Automated Scoring of the Pup Retrieval Test (MAS-PRT)

Seven days after the cessation of subcutaneous injections (vehicle, CBD, or THC), corresponding to postnatal day 5, dams underwent the pup retrieval test (PRT). The PRT is the most widely used assay for evaluating maternal care in fundamental and preclinical rodent studies and is frequently employed to study the impact of pharmacological and environmental interventions on maternal behavior (Smart, 1976; Zhang et al., 2019). In its standard form, the test measures the dam’s retrieval response to pup displacement from the nest, a task requiring a coordinated sequence of pup-directed sensorimotor behaviors elicited by the integration of multimodal pup stimuli (Winters et al., 2022, 2023).

Typical PRT protocols rely on manual scoring, which is prone to spatial and temporal inaccuracies. Plus, usually the experimenter that is annotating the test should be blind to the treatment. To address this limitation, we developed an automated detection system to extract PRT data, the Machine-Automated Scoring of the Pup Retrieval Test (MAS-PRT) to quantify maternal retrieval behavior at the level of individual pups and trials. The pipeline combines multi-animal DeepLabCut tracking of the dam and pups tracking (Figure 5A, C) with automated detection of the nest (Figure 5B), allowing extraction of first-encounter latency, full-retrieval latency, retrieval success, and maternal trajectories(Figure 5).

**Figure 5:**
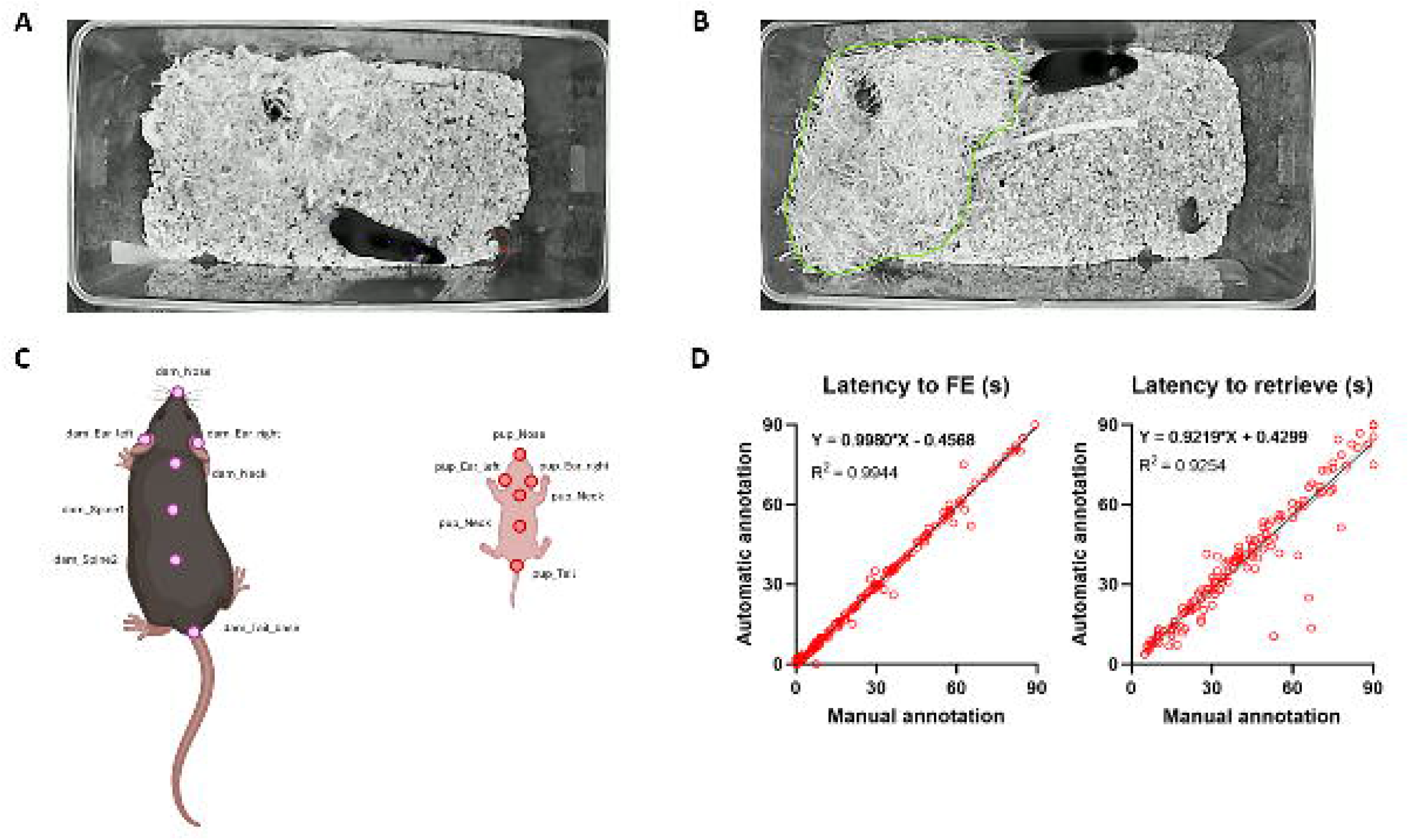
Validation of automatic pose and nest estimation. (A) Representative video frame captured during the test, showing DeepLabCut (DLC) tracking for both subjects (purple points: dam; red points: pup). (B) Example of automatic nest detection output generated using Detectron2. (C) Schematic illustration of the body parts annotated for DLC training. (C) Validation of the automated latency measurements against manual annotation. (D, left) Latency to first encounter (FE) and (D, right) latency to full retrieval were automatically detected and compared with manual annotations. The strong agreement between automated and manually annotated latency measurements demonstrates the accuracy of the automated event-detection approach. Regression resuls for (D, left) R^2^= 0.9944, p-value = <0.0001 and (D, right) R^2^= 0.9254, p-value = <0.0001. Data represented in XY plots, where each red point corresponds to an individual retrieval event of pups from all treatment conditions (n = 160 for A and 151 for B). Black lines represent the simple linear regression fit with 95% CI.

To assess measurement fidelity, automated measurements were compared with manual annotations performed by an experienced investigator blinded to treatment. Simple linear regression analyses revealed a high degree of concordance between the automated software and manual scoring for both behavioral metrics. Specifically, latency to first encounter (FE) exhibited a robust linear correlation with an *R*^2^ of 0.9944 (*p* < 0.0001; Figure 5D, left), while latency to full retrieval showed similarly strong agreement with an *R*^2^ of 0.9254 (*p* < 0.0001; Figure 5D, right). Mean deviations were 1.5% for first-encounter latency and 5.6% for retrieval latency. The tight clustering of data points along the regression line demonstrates minimal deviation and no systematic bias between human and automated measurement, confirming that the pipeline reliably and accurately captures maternal behavioral latencies.

These results establish that MAS-PRT can reproduce conventional latency measurements while providing automated access to additional pup-level and trajectory-based measures.

### Time-to-event analysis reveals enhanced encounter and retrieval probability of light-weight CBD-exposed pups

Pup retrieval is a widely used measure of parental responsiveness in rodents, primarily driven by auditory and olfactory cues (Kuroda & Tsuneoka, 2013; Okabe et al., 2013; Weber & Olsson, 2008; Winters et al., 2022). Although pups possess a non-functional auditory system at birth, they emit ultrasonic vocalizations (USVs) when isolated, which prompt dams to approach and retrieve them (Weber & Olsson, 2008). USVs emission serves as a proxy for pups affective state, and it is also linked to pup thermoregulation and body weight (Zimmer et al., 2019). Previous studies using the same prenatal CBD exposure protocol reported sex-specific differences in offspring outcomes, including USV patterns (Iezzi et al., 2022). Since pup calls are a primary motivator for retrieval (Ehret, 2005; Weber & Olsson, 2008), sex-related variations in vocalizations could influence maternal behavior. Additionally, litter size may impact retrieval performance, as larger litters require more attempts and are associated with smaller pup body weights (Parra-Vargas et al., 2020).

We next asked whether prenatal cannabinoid exposure altered maternal retrieval when pup-level variables and trial order were explicitly incorporated into the analysis. Because the 90 s test duration resulted in right-censored observations for pups that were not encountered or retrieved within the testing period, we used multivariate Cox proportional hazards models with robust standard errors to account for within-litter dependence. Models included treatment, pup sex, trial order, body weight, and treatment-by-weight interactions.

For first encounter, trial number was positively associated with the probability of encountering the displaced pup (HR = 1.096, p = 0.0029), consistent with progressive improvement across trials (Figure 6 A left panel; Table 2). Prenatal CBD exposure was associated with a significant treatment-by-body-weight interaction (HR = 0.161, p = 0.0009) (Figure 6B and Table 2), whereas the corresponding THC-by-weight interaction was not significant (HR = 1.234, p = 0.70) (Figure 6C and Table 2). Inspection of the model predictions indicated that the increased probability of encountering CBD-exposed pups was concentrated among pups with lower body weights, particularly below approximately 2.4 g at PND5. Pup sex was not significantly associated with first-encounter probability.

**Figure 6:**
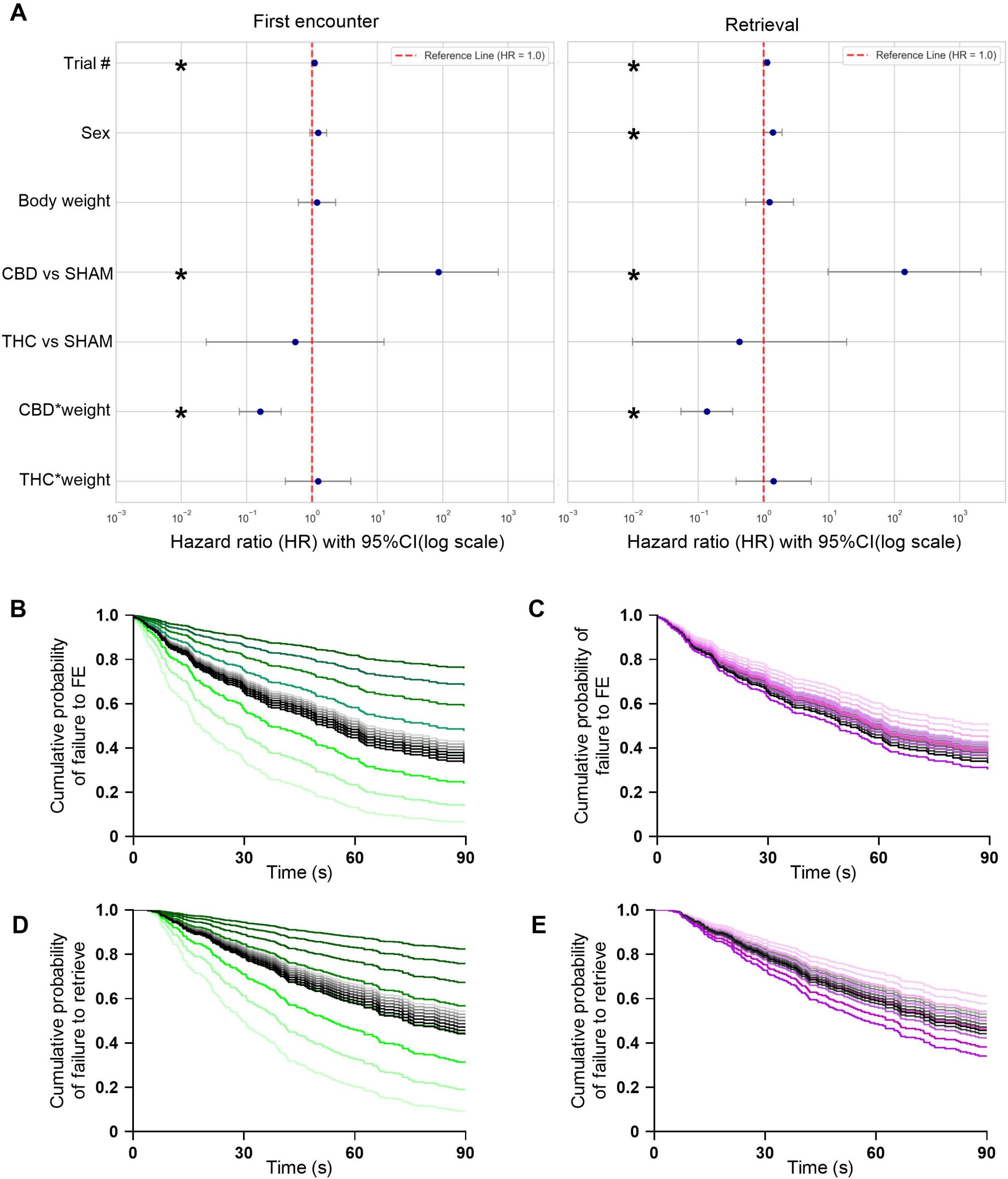
Maternal responses to CBD-exposed pups are modulated by pup body weight. (A) CBD treatment increases the likelihood of encounter and retrieve in a weight-dependent manner, where lighter pups are encountered and brought back to the nest more rapidly. Pup order in the test is a significant covariate, indicating a learning effect across trials. Forest plots displaying hazard ratios (HR) and 95% confidence intervals (CI) in logarithmic scale from multivariate Cox proportional hazards (Cox-PH) models; an HR>1 (with CI not touching 1) indicates an increased hazard (faster rate) of event completion. The regression model was run on the individual values for latency to retrieve of all of the mice in each litter, Cox regression included robust standard error to account for the litter effect (B-E) Predicted cumulative probability of task non-completion over time, modeled across a weight gradient (1.8-3.2 g in 0.2 g increments; shading intensity increases with weight). Color key: Black = Sham, Green = CBD, Purple = THC. (B) CBD treatment enhances encounter likelihood, moderated by body weight, but not (C) THC treatment. (D) CBD treatment enhances retrieval likelihood, moderated by body weight; (E) No weight-dependent effect is observed in the THC group, despite comparable growth restriction (see Figure 2). Analysis includes 82 control pups (13 litters), 100 CBD-exposed pups (13 litters), and 64 THC-exposed pups (8 litters).

**Table 2:**
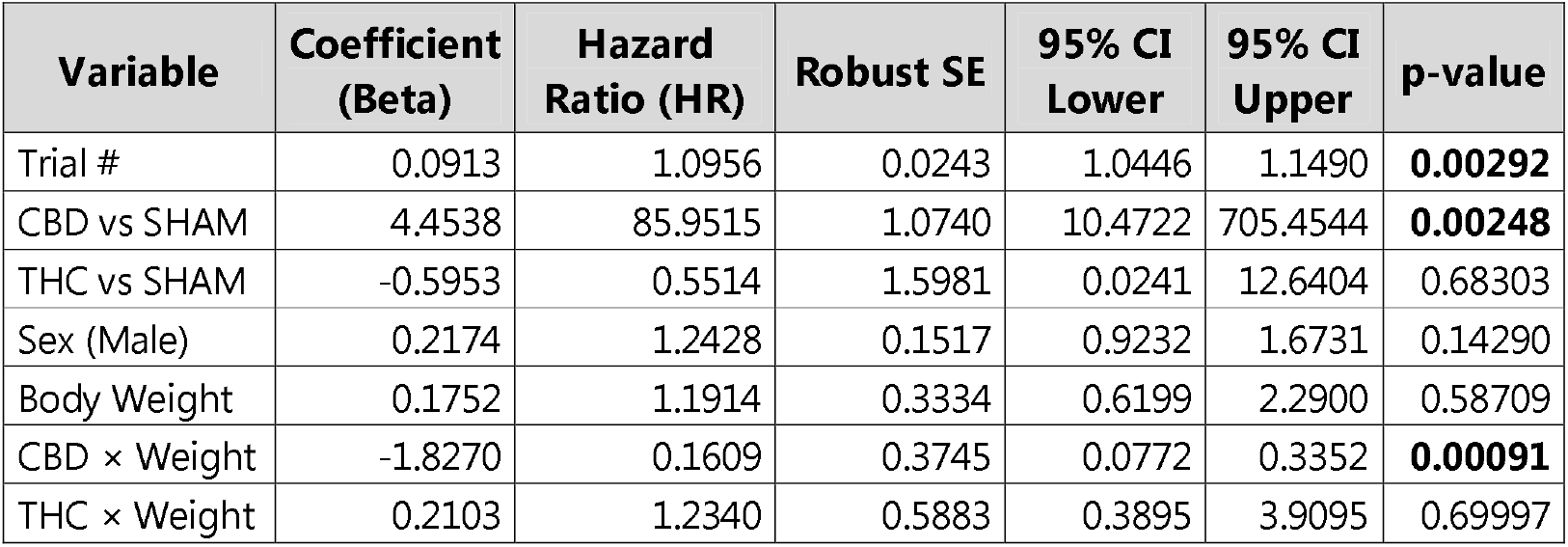
Cox Proportional Hazards regression analysis of latency to encounter the displaced pup. Results derived from multivariate Cox regression model. Hazard ratio (HR) indicates the relative risk of the event occurring compared to the reference level for each covariate. The time-to-event analysis highlights that CBD treatment increases the likelihood of first encounter, but this effect is strictly weight-dependent: only lightweight CBD pups have a higher chance of being encountered. Additionally, increasing trial numbers augment the chances of the event occurring (learning effect). Statistical significance set at *p*-value < 0.05.

| **Variable** | **Coefficient (Beta)** | **Hazard Ratio (HR)** | **Robust SE** | **95% CI Lower** | **95% CI Upper** | **p-value** |
| --- | --- | --- | --- | --- | --- | --- |
| Trial # | 0.0913 | 1.0956 | 0.0243 | 1.0446 | 1.1490 | **0.00292** |
| CBD vs SHAM | 4.4538 | 85.9515 | 1.0740 | 10.4722 | 705.4544 | **0.00248** |
| THC vs SHAM | -0.5953 | 0.5514 | 1.5981 | 0.0241 | 12.6404 | 0.68303 |
| Sex (Male) | 0.2174 | 1.2428 | 0.1517 | 0.9232 | 1.6731 | 0.14290 |
| Body Weight | 0.1752 | 1.1914 | 0.3334 | 0.6199 | 2.2900 | 0.58709 |
| CBD × Weight | -1.8270 | 0.1609 | 0.3745 | 0.0772 | 0.3352 | **0.00091** |
| THC × Weight | 0.2103 | 1.2340 | 0.5883 | 0.3895 | 3.9095 | 0.69997 |

The same analysis was applied to successful retrieval. Trial number again increased the probability of retrieval (HR = 1.123, p = 0.0002) (Figure 6 A right panel; Table 3). Prenatal CBD exposure was associated with a significant treatment-by-body-weight interaction (HR = 0.135, p = 0.0004) (Figure 6D, Table 3), whereas the corresponding THC interaction was not significant (HR = 1.421, p = 0.53) (Figure 6E, Table 3). Thus, the increased probability of retrieval following prenatal CBD exposure was conditional on pup body weight and was not observed as a general treatment effect across pups. Male sex was independently associated with increased retrieval probability in this model (HR = 1.379, p = 0.0338).

**Table 3:**
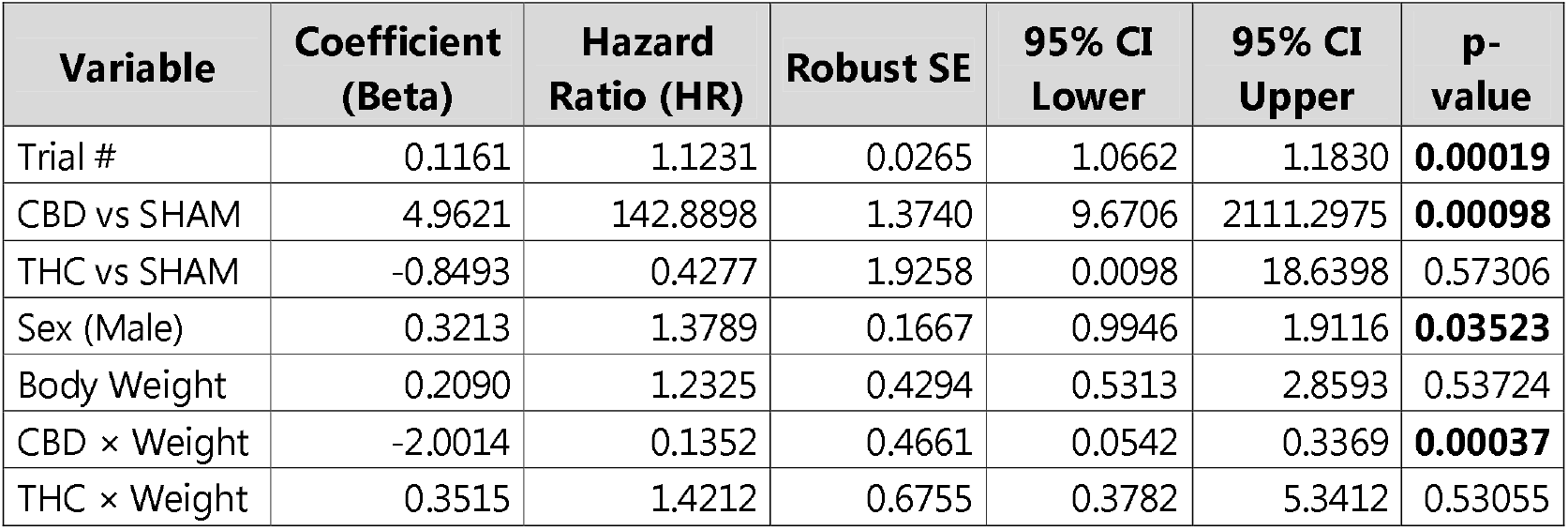
Cox Proportional Hazards regression analysis of successful pup retrieval. Results derived from multivariate Cox regression model. Hazard ratio (HR) indicates the relative risk of the event occurring compared to the reference level for each covariate. The time-to-event analysis highlights that CBD treatment increases the likelihood of retrieval, but this effect is strictly weight-dependent: only lightweight CBD pups have a higher chance of being retrieved. The probability of the event is positively associated with both trial frequency and male biological sex. Statistical significance set at *p*-value < 0.05.

Animals were randomly selected to perform the test, but to ensure that the selection was not biased by pup size, Levene’s tests for homogeneity of variance were performed. No significant differences in body weight variance were found across trial numbers, neither in the general population (p = 0.798) nor when analyzed within specific treatment groups (Sham p= 0.981; CBD p= 0.596; THC = 0.994), confirming that pup selection was random with respect to weight (Supplementary Table 1).

### Prenatal CBD and THC exposure alter retrieval trajectories for female offspring

Because MAS-PRT also provides continuous measures of maternal movement, we next examined the trajectory used to reach and retrieve each pup (Figure 7). Neither prenatal CBD nor THC exposure altered the distance traveled before the first encounter or the distance traveled between first encounter and retrieval. This was evidenced by the distance covered to reach the pup (Figure 7A), the distance traveled between the first encounter and retrieval (Figure 7B). However, total distance traveled during the complete retrieval sequence was reduced when dams retrieved female offspring from both prenatal exposure groups (Figure 7C).

**Figure 7:**
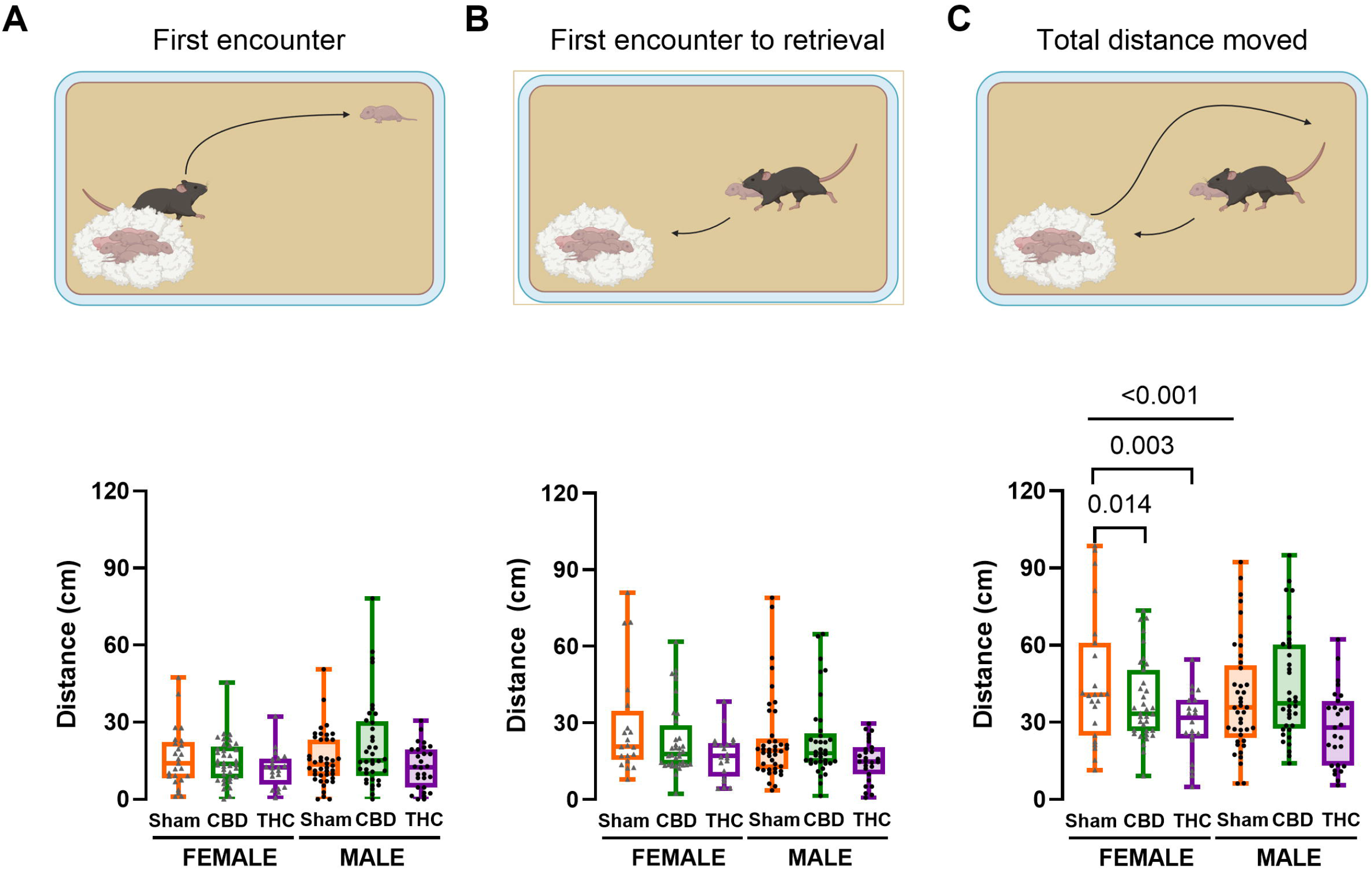
Dams implement shorter trajectories to retrieve CBD- and THC-exposed female progeny. (A-C) Quantification of total distance moved across groups. There were no significant differences in the (A) distance traveled to locate pups and (B) distance covered during pup retrieval, but the (C) total distance traveled during the complete retrieval sequence. Data are presented as box-and-whisker plots showing median, minimum, and maximum values; each data point represents the mean performance per dam across all trials. Analysis includes 82 control pups (13 litters), 100 CBD-exposed pups (13 litters), and 64 THC-exposed pups (8 litters). Statistical analysis was performed using the linear mixed models with Litter ID as a random intercept were used to account for litter-level variance. Post-hoc pairwise comparisons were adjusted for multiple testing across the 7 planned contrasts using the Benjamini–Hochberg False Discovery Rate (FDR) procedure, with significance defined as *p* < 0.05.

Thus, prenatal CBD and THC exposure did not produce a generalized alteration in maternal locomotion during the retrieval task. Instead, both exposures were associated with a more spatially constrained retrieval trajectory specifically for female offspring (see statistical details in Table 4).

**Table 4:**
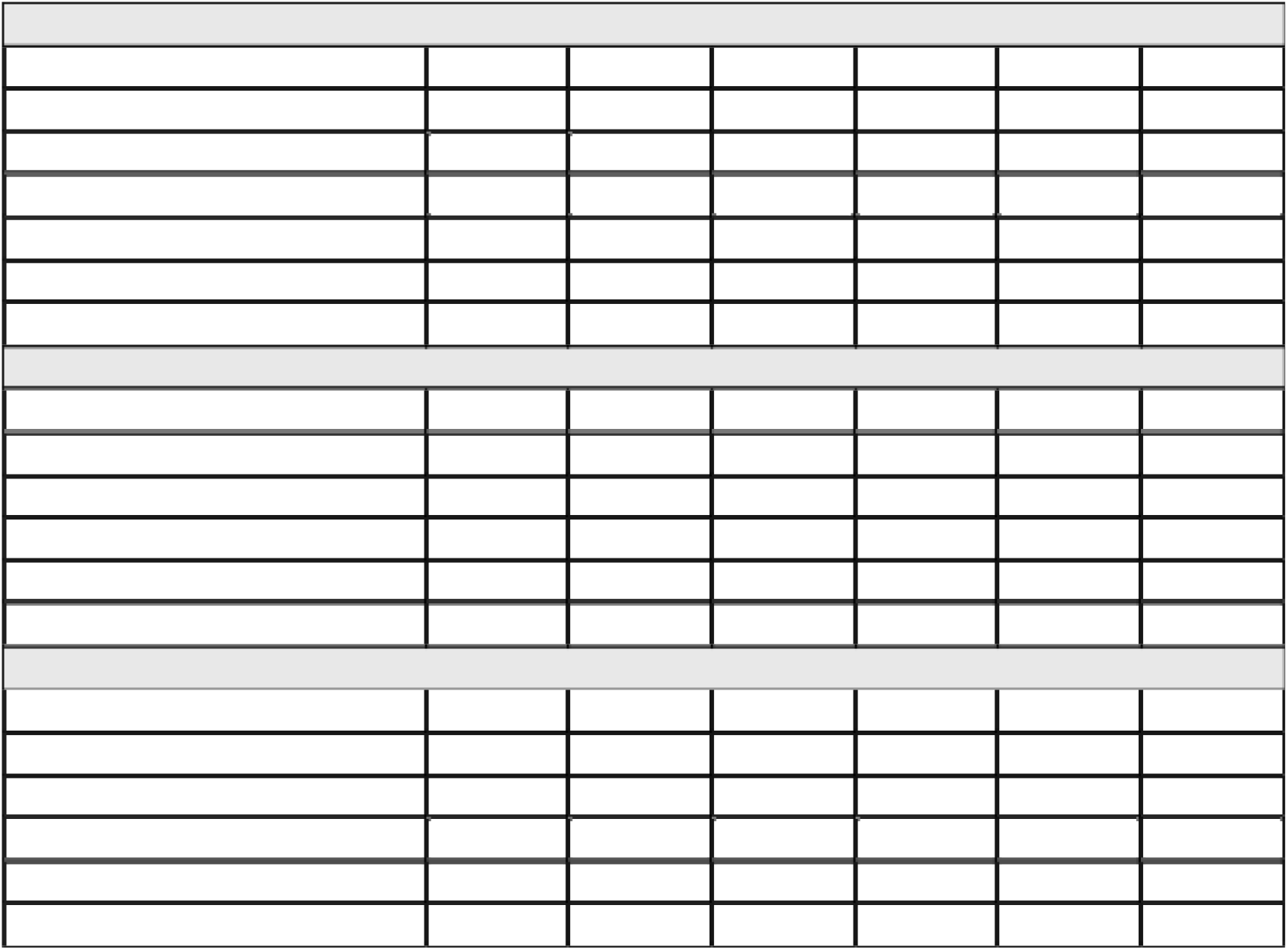
Linear mixed-effects model results dam’s distance moved during retrieval behavior. Fixed-effects estimates, standard errors, z-scores, p-values, and 95% confidence intervals derived from full factorial LMMs (Metric ∼ Sex * Treatment) with Litter ID included as a random intercept term. Statistical significance: p< 0.05. (A) Fixed factors accounted for a modest proportion of variance 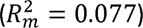, whereas the conditional model accounted for 23.18% 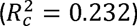. This highlights that exploratory search prior to pup contact is heavily influenced by litter-level variance 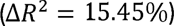 rather than pup sex or drug treatment. (B) Fixed factors accounted for 4.13% of variance 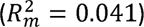, with total explained variance rising to 8.19% when including litter identity 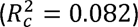. (C) The overall model explained 9.04% of total variance 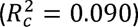, with fixed experimental factors accounting for 8.64% 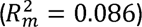. The small difference between 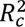 and 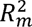 indicates minimal litter-level clustering during total retrieval.

### Differential learning curves across treatments

Finally, we examined whether differences in maternal experience across successive retrieval trials could account for the weight-dependent CBD phenotype. By looking at the evolution of individual time taken by each dam to first encounter (Figure 8A, Table 5), and retrieve (Figure 8B, Table 5) each mouse across trials, Sham dams display clearly shorter latencies as the trials advance.

**Figure 8:**
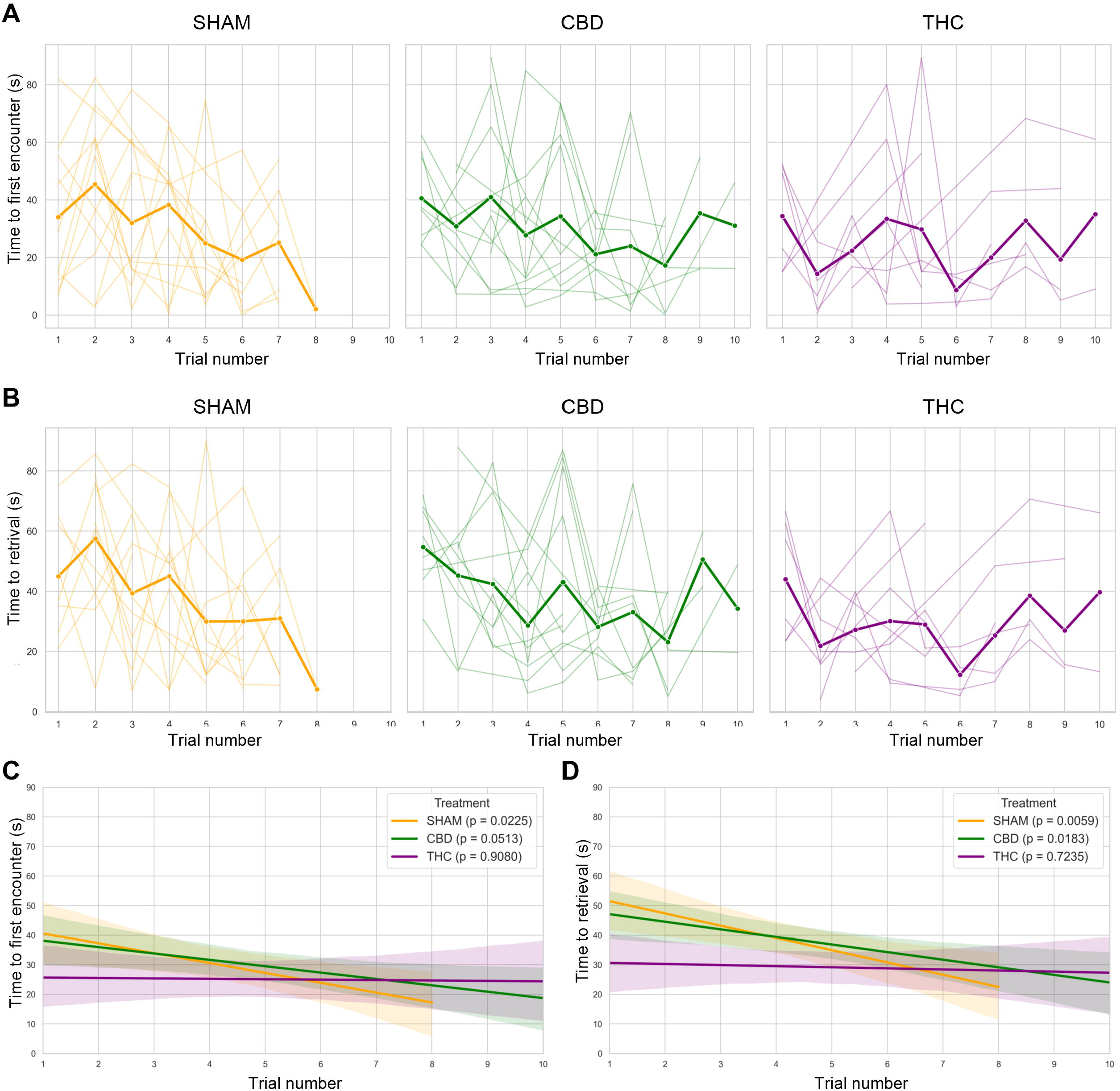
Learning curves across trials. (A) Time to first encounter across trial numbers, showing individual subject trajectories (thin lines) alongside group mean trends (thick lines) for SHAM (orange), CBD (green), and THC (purple) groups. (B) Time to retrieval across trial numbers, displaying dam individual performance progression and group means for each respective treatment group. (C) Linear regression fits with 95% confidence intervals depicting the trend in time to first encounter across trials for each treatment group, including group-specific p-values (SHAM: p= 0.0225; CBD: p=0.0513; THC: p=0.9080). (D) Linear regression fits with 95% confidence intervals showing the trend in time to retrieval across trials, with corresponding group-specific p-values (SHAM: p=0.0059; CBD: p=0.0183; THC: p=0.7235).

**Table 5:**
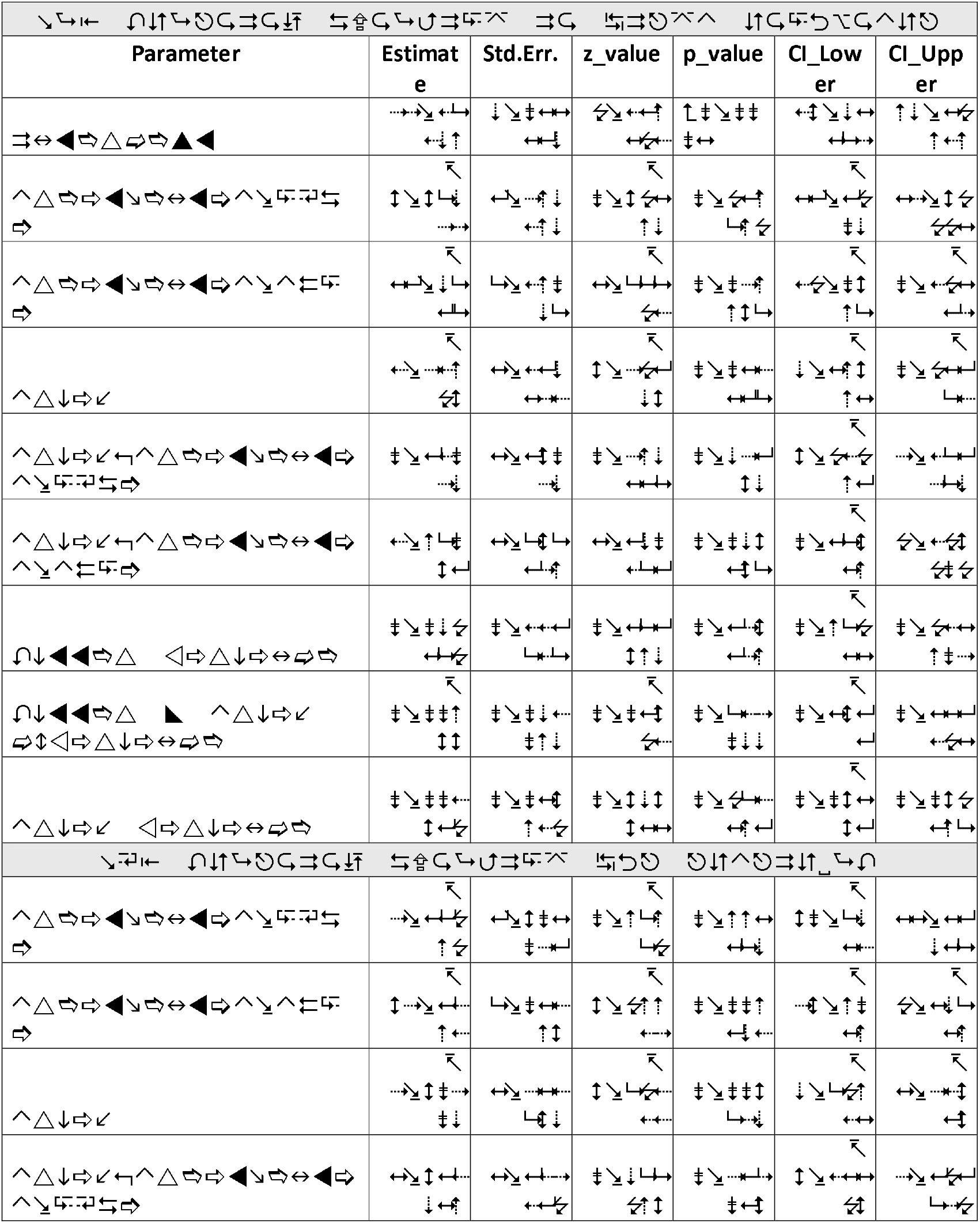
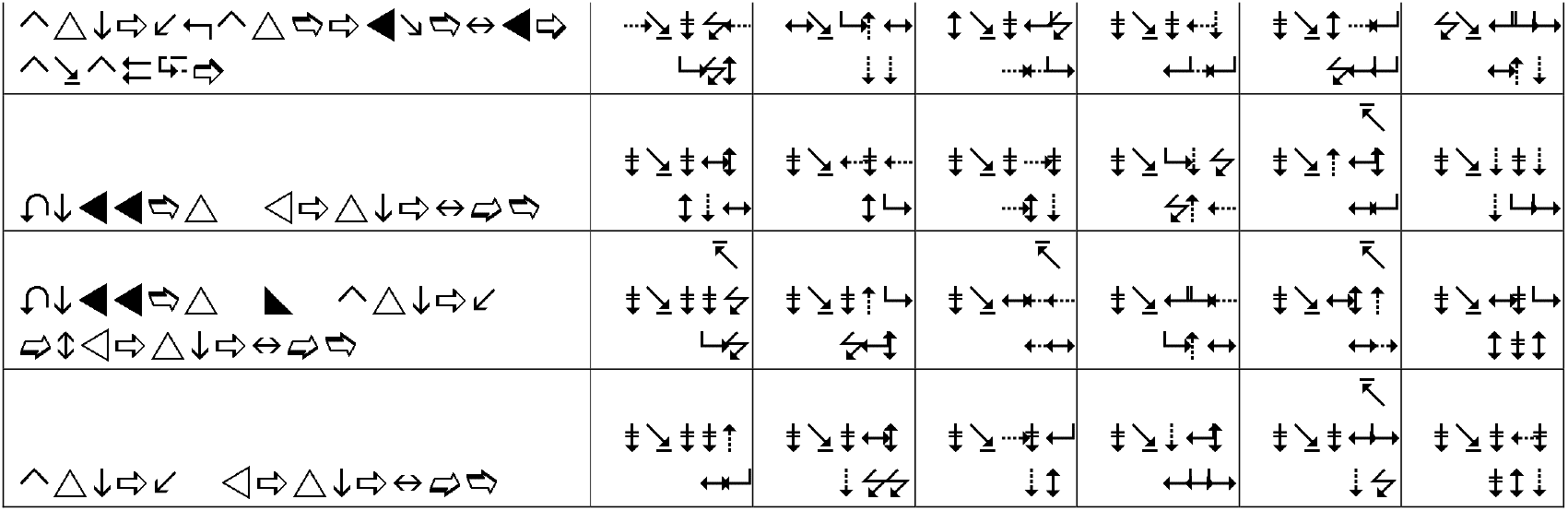
Linear mixed-effects model results on learning curve along trials. (A) Fixed factors (Treatment, Trial, and Treatment ×Trial interactions) account for 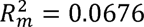 (6.76%) of total variance. The inclusion of random litter intercepts 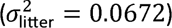 and trial slopes 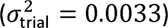 increases total explained variance to 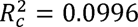 (9.96%). (B) Fixed factors explain 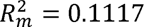 (11.17%) of behavioral variance. Combined fixed and random variance components account for 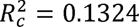 (13.24%) of total variance, reflecting an additional 2.07% contribution from litter identity 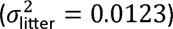.

First, we fitted the individual data to linear regressions to confirm whether, within each treatment, increased trial number would lead to shorter latencies to perform the task. As trials advanced Sham dams got a faster performance in the task, for both, encounter (Figure 8C) and full retrieval (Figure 8D). Similar trends were observed for CBD-treated dams, which showed significant improvement in retrieval time, alongside a near-significant trend for encounter latency.

Due to the nested nature of the data, we performed linear mixed models (Table 5) to account for the litter effect in the analysis, because across trials the dam would stop being experimentally naïve. In addition, this analysis allowed us to correct for the fact that not all litter are the same size, thus dams had different number of trials to learn. The analysis for first encounter latency revealed a significant overall main effect of trial number (*β* = −3.44, SE = 1.39, *z* = −2.48, *p* = 0.013), indicating that Sham subjects successfully reduced their latency to make a first encounter as training progressed. While CBD-treated animals did not significantly differ from SHAM controls at trial 1 (*β* = −2.30, SE = 8.46, *z* = −0.27, *p* = 0.786) or in their learning rate (*β* = +0.83, SE = 1.82, *z* = 0.46, *p* = 0.648), THC-treated animals exhibited a significant baseline difference, starting with a substantially lower first encounter latency on trial 1 compared to SHAM controls (*β* = −18.70, SE = 9.35, *z* = −2.00, *p* = 0.046). The interaction between trial and THC treatment was not proven, but close to statistical significance (*β* = +3.59, SE = 1.93, *z* = 1.86, *p* = 0.063), suggesting that most likely the decreased learning curve steepness was due to THC dams taking shorter time to complete the test from the very beginning.

Similarly, analysis of time to retrieval (Table 5) showed a robust main effect of Trial in the SHAM control group (*β* = −4.20, SE = 1.41, *z* = −2.97, *p* = 0.003), reflecting learning and performance improvement over trials. Baseline performance (Trial 1) did not differ significantly between CBD and SHAM groups (*β* = −4.89, SE = 8.20, *z* = −0.60, *p* = 0.551). However, THC-treated subjects demonstrated a highly significant reduction in initial retrieval latency on the first trial relative to SHAM controls (*β* = −24.84, SE = 9.01, *z* = −2.76, *p* = 0.006). Furthermore, the interaction term for trial by THC treatment was statistically significant (*β* = +4.07, SE = 1.95, *z* = 2.09, *p* = 0.037). These findings indicate that the weight-dependent retrieval phenotype observed following prenatal CBD exposure cannot be explained by an altered acquisition of the retrieval task. Instead, CBD-exposed dams showed trial-to-trial dynamics comparable to controls, whereas prenatal THC exposure was associated with a distinct pattern of initially rapid retrieval and reduced subsequent improvement.

## DISCUSSION

The present study investigated maternal behavior following prenatal CBD and THC exposure using conventional measures together with automated, pup-level analysis of the pup retrieval test. Three findings emerge. First, prenatal CBD or THC exposure did not produce a generalized disruption of gestational outcomes, nest building, or daily nursing behavior under the conditions tested. Second, automated time-to-event analysis revealed a selective weight-dependent alteration in maternal responsiveness following prenatal CBD exposure: lighter CBD-exposed pups were encountered and retrieved with a higher probability than predicted from their treatment and body weight alone. Third, prenatal CBD and THC produced distinct alterations in retrieval trajectories and trial-by-trial performance. Together, these findings illustrate how pup-level analysis can reveal features of maternal behavior that are not captured by conventional mean dam performance.

The most prominent finding was the interaction between prenatal CBD exposure and pup body weight in predicting both first encounter and successful retrieval. Importantly, this was not a generalized acceleration of retrieval by CBD-exposed dams. Rather, the association emerged as a function of pup weight, with the effect concentrated among lower-weight pups. This distinction is important because both CBD- and THC-exposed pups showed reduced body weight at PND5, yet only CBD exposure was associated with a significant treatment-by-weight interaction in the retrieval analyses. The maternal behavioral phenotype therefore cannot be explained simply by the presence of smaller pups.

Several mechanisms could contribute to this selective association. Pup body weight is correlated with developmental state and can influence the sensory cues available to the dam, including ultrasonic vocalizations and other pup-derived signals (Ehret, 2005; Okabe et al., 2013; Zimmer et al., 2019). Previous work using the same prenatal CBD exposure paradigm identified sex-dependent alterations in early-life ultrasonic vocalizations (Iezzi et al., 2022), raising the possibility that prenatal CBD exposure modifies the signals emitted by offspring and thereby changes maternal responsiveness. Alternatively, prenatal CBD exposure could alter sensory processing or sensorimotor integration in the dam, changing the salience assigned to cues associated with smaller offspring. The present experiments cannot distinguish between these possibilities because pup vocalizations were not recorded during the retrieval task.

The behavioral phenotype is therefore best interpreted as an alteration in the maternal responsiveness rather than as a global increase in maternal motivation. The preservation of nest building and daily nursing behavior, together with the absence of a generalized reduction in retrieval performance, argues against a broad disruption of maternal care. Instead, prenatal CBD exposure appears to modify how maternal retrieval behavior is expressed in relation to characteristics of individual offspring.

A central contribution of this study is methodological as well as biological. Conventional PRT displaces several pups at the same time, which difficult the establishment of relationships between maternal behavior and properties of individual pups or the dam’s experience during successive trials (Hahn & Lavooy, 2005; Weber & Olsson, 2008). MAS-PRT allowed us to link each behavioral event to an individual pup, its body weight and sex, its position relative to the nest, and its position within the sequence of retrieval trials. The resulting Cox proportional hazards analysis was particularly useful because pups that were not encountered or retrieved within the 90 s test period were treated as censored observations rather than simply excluded or assigned an arbitrary latency. Incorporating pup weight, sex, treatment, trial order, and treatment-by-weight interactions therefore allowed us to test conditional effects that would be difficult to resolve using a conventional group-level comparison.

This analytical perspective provides an important interpretation of the present findings. The absence of a strong treatment effect in conventional latency measures already reported for CBD progeny (Compagno et al., 2025) does not necessarily indicate that maternal retrieval is unaffected. Instead, a treatment effect may emerge only when behavioral responses are examined in relation to characteristics of the pup. In this case, the CBD-associated phenotype became apparent only after pup body weight was explicitly modeled.

The automated trajectory analysis provided a complementary dimension. Neither CBD nor THC altered the distances traveled to reach the pup or from first encounter to retrieval, arguing against a generalized locomotor alteration during the task. However, both exposures were associated with shorter complete retrieval trajectories for female offspring. This result illustrates an additional advantage of automated tracking: changes in the spatial organization of behavior can be detected even when conventional temporal measures remain broadly similar. We interpret these differences as altered retrieval kinematics rather than evidence for greater retrieval efficiency, since the present study did not directly test the functional consequences of trajectory length.

In addition, we reported that CBD and THC dams take shorter trajectories to retrieve their female offspring, even though this does not end on THC female offspring having higher likelihood to be retrieved. The differential weight-dependent reactivity of CBD dams cannot be explained by faster learning acquisition of the task, as show learning patterns comparable to Sham dams.

Our findings complement previous studies reporting relatively preserved maternal behavior following prenatal THC exposure (Bara et al., 2018; Frau et al., 2019, 2019). They also extend recent work examining the consequences of prenatal CBD exposure, which has demonstrated effects on offspring development without necessarily producing an overt maternal-care deficit (Compagno et al., 2025). Differences in exposure route, dose, timing, and PRT design make direct comparisons between studies difficult. Nevertheless, the present results suggest that preserved conventional measures of maternal behavior do not exclude more selective changes in the mother-offspring interaction.

Regarding our automated analysis to annotate PRT, unlike Simba-based approaches like BAMBI (Winters et al., 2023) that often require manual nest (ROIs) delimitation, MAS-PRT offers a fully automated, batch-processable workflow. Despite these advances, several limitations remain. First, pup ultrasonic vocalizations were not recorded simultaneously with retrieval behavior. Because prenatal CBD exposure has previously been associated with altered early-life vocalization patterns (Iezzi et al., 2022), simultaneous measurement of pup acoustic signals and maternal behavior will be important for determining whether altered offspring signaling contributes to the present phenotype. Second, the present analysis was performed using top-view recordings in C57BL/6J mice; application of MAS-PRT to other strains, recording configurations, or behavioral paradigms may require additional model training or validation. Third, the PRT provides a relatively brief assay of maternal responsiveness. Although the trial-by-trial analysis provides evidence for experience-dependent changes during the test, additional paradigms will be required to determine whether prenatal cannabinoid exposure affects broader aspects of maternal learning or behavioral flexibility.

Finally, the present findings should not be interpreted as evidence that prenatal CBD or THC exposure is without developmental consequences. The absence of a generalized maternal-care deficit occurred alongside a measurable reduction in early postnatal pup body weight and, in the case of CBD, a selective alteration in maternal retrieval dynamics. Prenatal cannabinoid exposure has been associated with a range of developmental effects across species and experimental paradigms (Higuera-Matas et al., 2015; Jenkins et al., 2025; Scheyer et al., 2019). The present study adds a specific behavioral dimension by showing that the maternal response to individual offspring can be altered without producing a global impairment in maternal care.

## Author Contributions

A.C.-R.: conceptualization, data curation, formal analysis, validation, writing—review and editing. B.S.: formal analysis, validation. J.F.: formal analysis, validation. J.P.: formal analysis, validation. D.I.: data curation. P.C.: conceptualization, supervision. O.J.J.M.: conceptualization, supervision, funding acquisition, methodology, project administration, writing—original draft, review, and editing. All authors have read and agreed to the published version of the manuscript.

## Acknowledgements

The authors are grateful to the Chavis-Manzoni team members for helpful discussions.

---

**Supplementary Figure 1:**
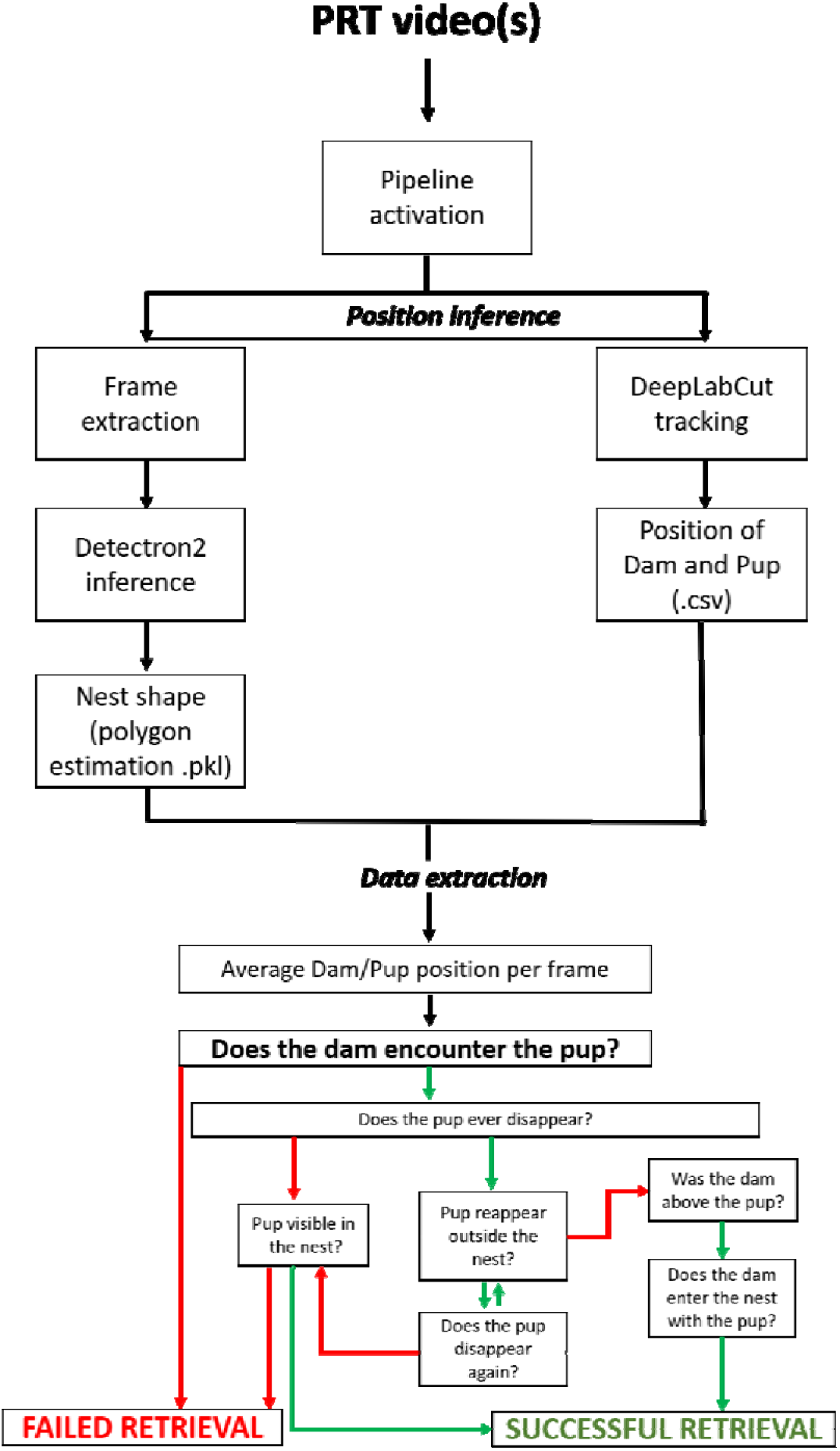
Schema of the pipeline required for the automated analysis. Once the video is uploaded in the software, two different position extractions are performed: nest area detection and two-animal tracking. The positional data is used to determine whether the dam has encountered the pup and the frame in which it has been retrieved.

**Supplementary Figure 2:**
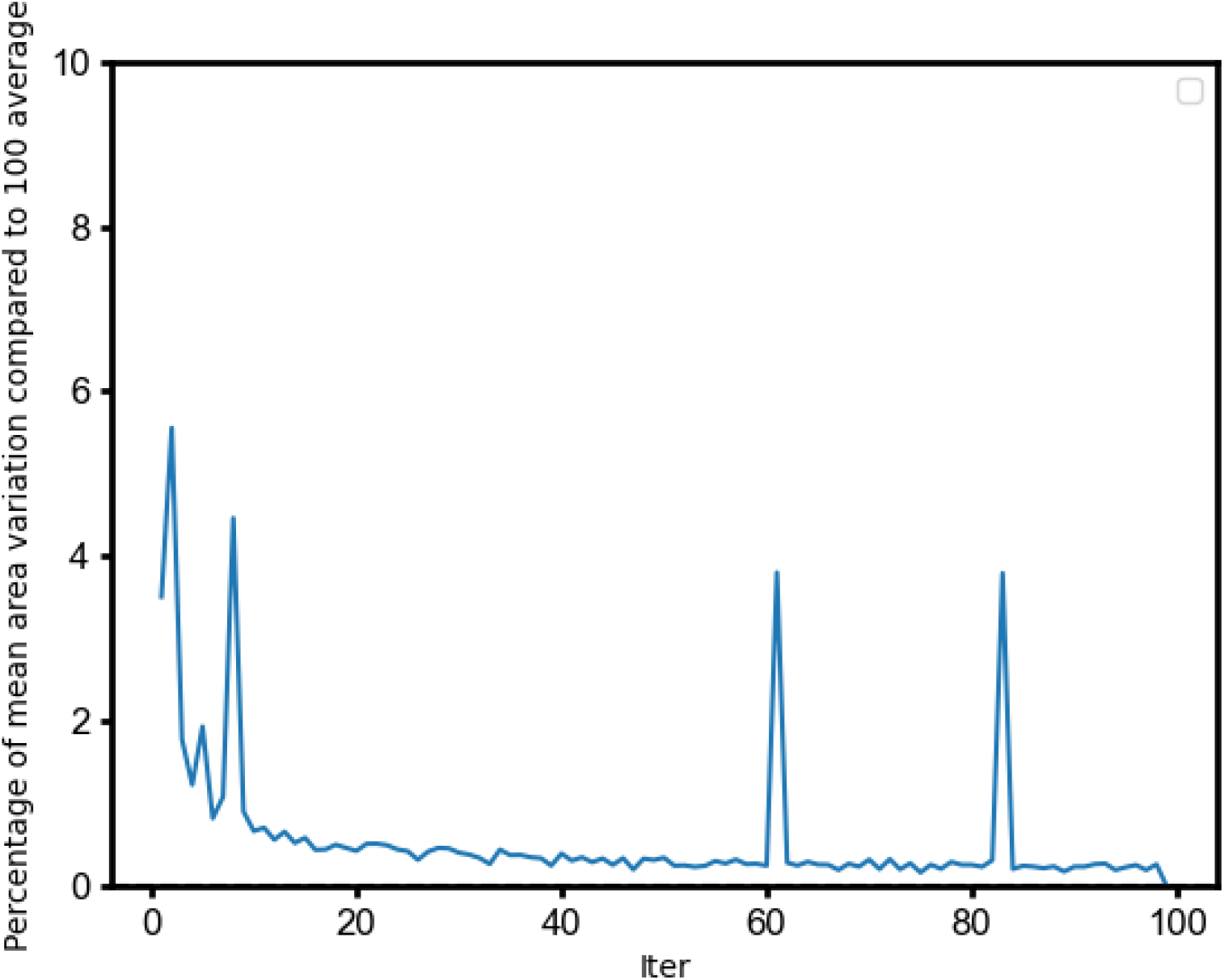
**Effect on accuracy detection of number of frames used to estimate nest area**. This analysis evaluates how the number of video frames used to draw the nest mask influences the accuracy of nest area estimation. A reference value was established by averaging measurements across 100 frames. The graph plots the percentage difference between estimates obtained with fewer frames and this reference value. The X-axis shows the number of frames included in the calculation, while the Y-axis shows the deviation (%) from the reference area. As the number of frames increases, the estimates converge toward the reference, indicating that larger frame samples improve stability and accuracy; however, 20 frames were sufficient for correct estimation.

**Supplementary Figure 3:**
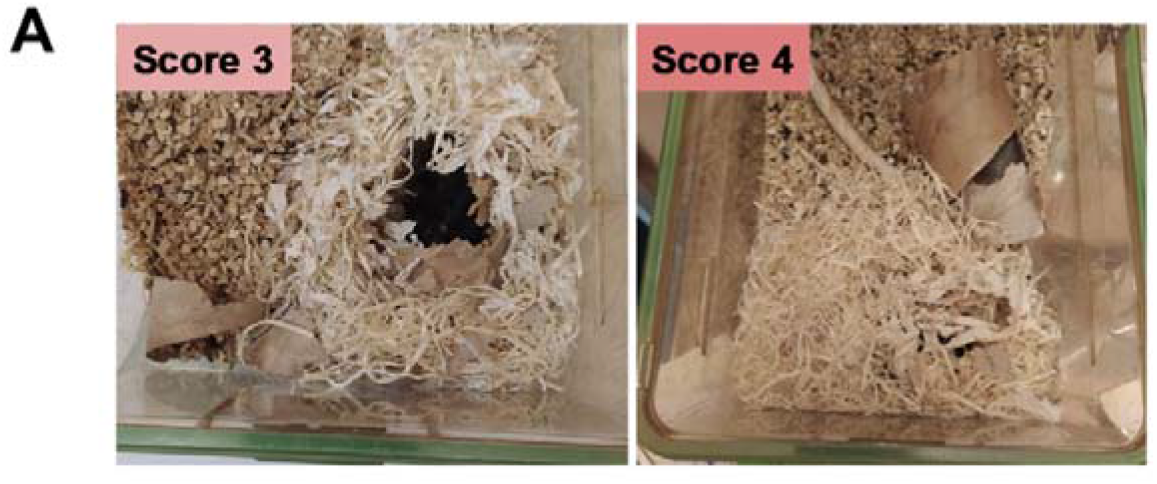
Home-cage nest grading. (A) Representative images illustrate the two most frequent nest quality scores: Score 3 corresponds to fully shredded nesting material forming walls tall enough to cover the dam; Score 4 reflects a complete enclosure with a roof, fully covering the dam.

**Supplementary Figure 4:**
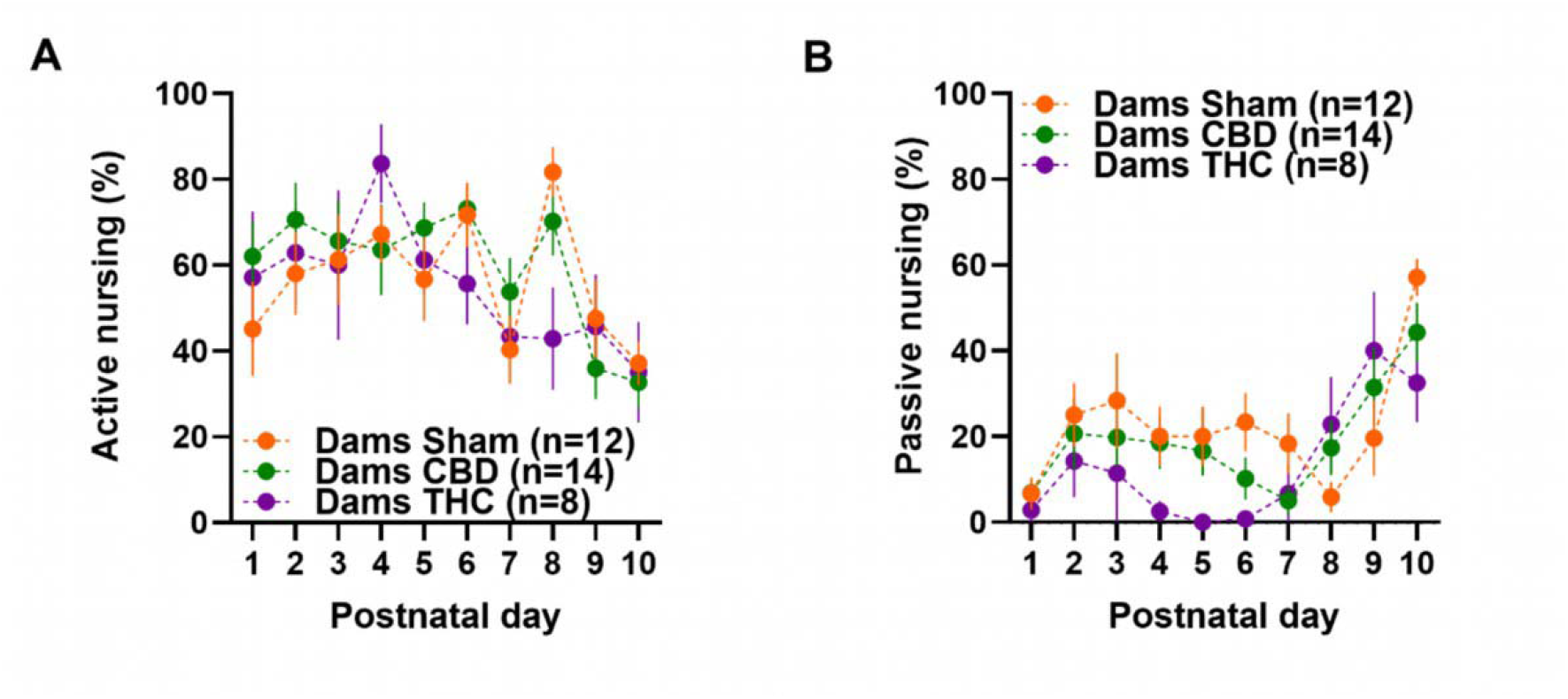
Nursing patterns during the first 10 postnatal days. Nursing style shifts over time. Specifically, (A) active nursing predominates in the early postpartum period and transitions to (B) passive nursing by the end of the follow-up (PND 9–10). (A–B) Time-course data are presented as mean ± SEM in XY plots. Group sizes: Sham (n = 12), CBD (n = 14), THC (n = 8). Statistical analysis was performed via mixed-effects models with the Geisser-Greenhouse correction, followed by Tukey’s post hoc multiple comparison test. Only results with *p* < 0.05 were considered statistically significant and are reported.

**Supplementary Table 1:**
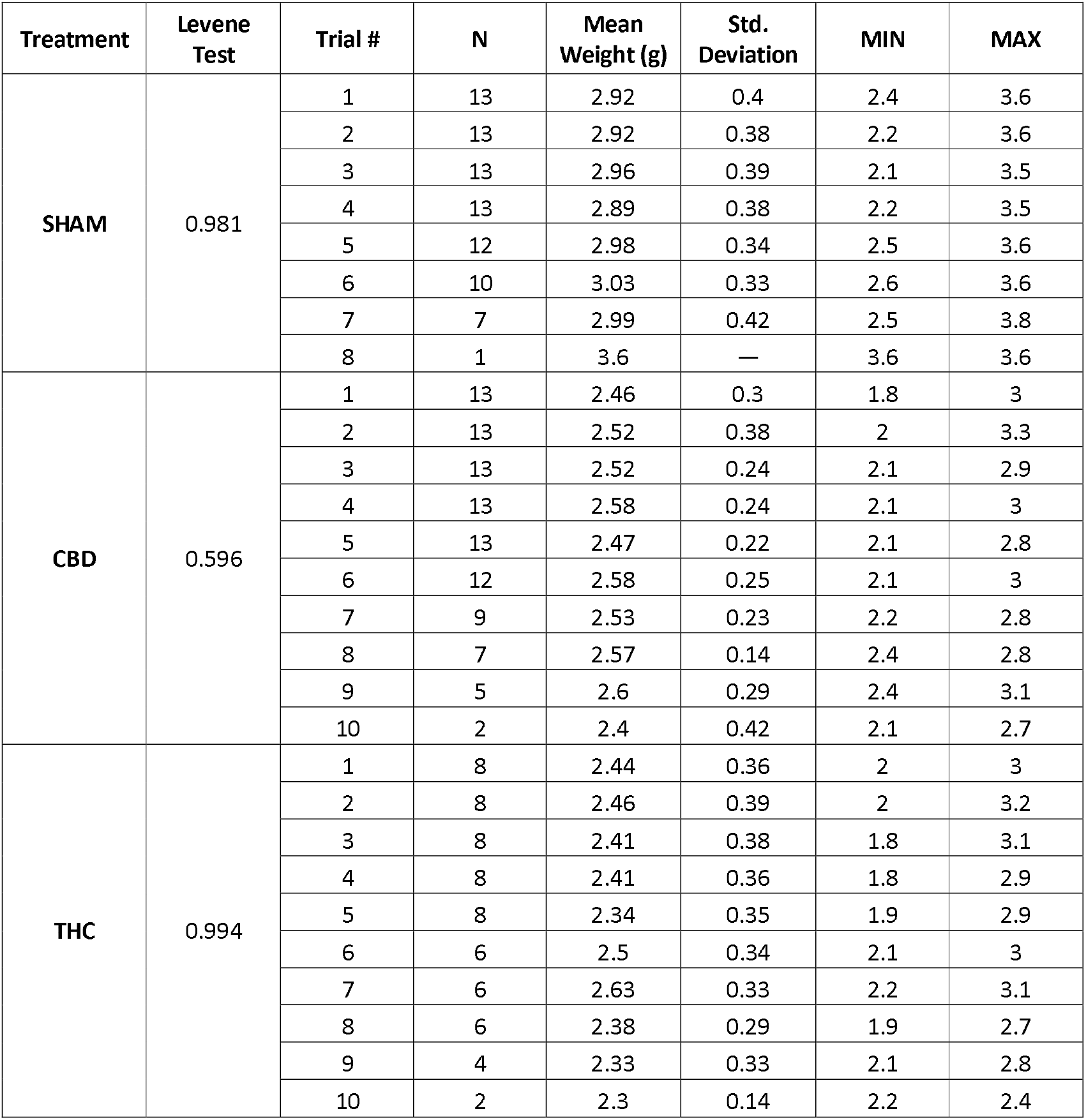
The random selection of pup order for the PRT task is not biased by body weight. Levene’s test indicated no significant differences in weight variability based on the order in which pups from the same litter were tested. The table summarizes pup weight statistics for each treatment group, organized by testing order. Reported values include treatment group, Levene’s test *p*-value, testing order, number of pups (n), mean pup weight (g), standard deviation (std deviation), and minimum (MIN) and maximum (MAX) pup weights.

## REFERENCES

Bailey, S., & Isogai, Y. (2022). Parenting as a model for behavioural switches. Current Opinion in Neurobiology, 73, 102543. 10.1016/j.conb.2022.102543

Bara, A., Manduca, A., Bernabeu, A., Borsoi, M., Serviado, M., Lassalle, O., Murphy, M., Wager-Miller, J., Mackie, K., Pelissier-Alicot, A.-L., Trezza, V., & Manzoni, O. J. (2018). Sex-dependent effects of in utero cannabinoid exposure on cortical function. eLife, 7, e36234. 10.7554/eLife.36234

Bhatia, D., Battula, S., Mikulich-Gilbertson, S., Sakai, J., & Hammond, D. (2024). Cannabidiol-Only Product Use in Pregnancy in the United States and Canada: Findings From the International Cannabis Policy Study. Obstetrics & Gynecology, 144(2), 156–159. 10.1097/AOG.0000000000005603

Bowers, J. M., Perez-Pouchoulen, M., Edwards, N. S., & McCarthy, M. M. (2013). Foxp2 Mediates Sex Differences in Ultrasonic Vocalization by Rat Pups and Directs Order of Maternal Retrieval. The Journal of Neuroscience, 33(8), 3276–3283. 10.1523/JNEUROSCI.0425-12.2013

Bridges, R. S. (2015). Neuroendocrine regulation of maternal behavior. Frontiers in Neuroendocrinology, 36, 178–196. 10.1016/j.yfrne.2014.11.007

Calapai, F., Cardia, L., Calapai, G., Di Mauro, D., Trimarchi, F., Ammendolia, I., & Mannucci, C. (2022). Effects of Cannabidiol on Locomotor Activity. Life, 12(5), 652. 10.3390/life12050652

Castel, P., Barbier, M., Poumerol, E., Mandon-Pépin, B., Tassistro, V., Lepidi, H., Pelissier-Alicot, A.-L., Manzoni, O. J., & Courbiere, B. (2020). Prenatal cannabinoid exposure alters the ovarian reserve in adult offspring of rats. Archives of Toxicology, 94(12), 4131–4141. 10.1007/s00204-020-02877-1

Cataldo, I., Azhari, A., Coppola, A., Bornstein, M. H., & Esposito, G. (2019). The Influences of Drug Abuse on Mother-Infant Interaction Through the Lens of the Biopsychosocial Model of Health and Illness: A Review. Frontiers in Public Health, 7, 45. 10.3389/fpubh.2019.00045

Compagno, M. K., Silver, C. R., Cox-Holmes, A., Basso, K. B., Bishop, C., Bernstein, A. M., Carley, A., Cazorla, J., Claydon, J., Crane, A., Crespi, C., Curley, E., Dolezel, T., Franck, E., Heiden, K., Huffstetler, C. M., Loeven, A. M., May, C. A., Maykut, N., … Fadool, D. A. (2025). Maternal ingestion of cannabidiol (CBD) in mice leads to sex-dependent changes in memory, anxiety, and metabolism in the adult offspring, and causes a decrease in survival to weaning age. Pharmacology Biochemistry and Behavior, 247, 173902. 10.1016/j.pbb.2024.173902

Curley, J. P., & Champagne, F. A. (2016). Influence of Maternal Care on the Developing Brain: Mechanisms, Temporal Dynamics and Sensitive Periods. Frontiers in neuroendocrinology, 40, 52–66. 10.1016/j.yfrne.2015.11.001

Deacon, R. M. J. (2006). Assessing nest building in mice. Nature Protocols, 1(3), 1117–1119. 10.1038/nprot.2006.170

DeVuono, M. V., Nashed, M. G., Sarikahya, M. H., Kocsis, A., Lee, K., Vanin, S. R., Hudson, R., Lonnee, E. P., Rushlow, W. J., Hardy, D. B., & Laviolette, S. R. (2024). Prenatal tetrahydrocannabinol and cannabidiol exposure produce sex-specific pathophysiological phenotypes in the adolescent prefrontal cortex and hippocampus. Neurobiology of Disease, 199, 106588. 10.1016/j.nbd.2024.106588

Ehret, G. (2005). Infant Rodent Ultrasounds—A Gate to the Understanding of Sound Communication. Behavior Genetics, 35(1), 19–29. 10.1007/s10519-004-0853-8

El Marroun, Tiemeier, Steegers, Jaddoe, & Et., A. (2009). Intrauterine Cannabis Exposure Affects Fetal Growth Trajectories: The Generation R Study. Journal of the American Academy of Child & Adolescent Psychiatry. 10.1097/chi.0b013e3181bfa8ee

Frau, R., Miczán, V., Traccis, F., Aroni, S., Pongor, C. I., Saba, P., Serra, V., Sagheddu, C., Fanni, S., Congiu, M., Devoto, P., Cheer, J. F., Katona, I., & Melis, M. (2019). Prenatal THC exposure produces a hyperdopaminergic phenotype rescued by pregnenolone. Nature Neuroscience, 22(12), 1975–1985. 10.1038/s41593-019-0512-2

Gammie, S. C. (2005). Current models and future directions for understanding the neural circuitries of maternal behaviors in rodents. Behavioral and Cognitive Neuroscience Reviews, 4(2), 119–135. 10.1177/1534582305281086

Hahn, M. E., & Lavooy, M. J. (2005). A review of the methods of studies on infant ultrasound production and maternal retrieval in small rodents. Behavior Genetics, 35(1), 31–52. 10.1007/s10519-004-0854-7

Harkany, T., Guzmán, M., Galve-Roperh, I., Berghuis, P., Devi, L. A., & Mackie, K. (2007). The emerging functions of endocannabinoid signaling during CNS development. Trends in Pharmacological Sciences, 28(2), 83–92. 10.1016/j.tips.2006.12.004

Hess, S. E., Rohr, S., Dufour, B. D., Gaskill, B. N., Pajor, E. A., & Garner, J. P. (2008). Home improvement: C57BL/6J mice given more naturalistic nesting materials build better nests. Journal of the American Association for Laboratory Animal Science: JAALAS, 47(6), 25–31.

Higuera-Matas, A., Ucha, M., & Ambrosio, E. (2015). Long-term consequences of perinatal and adolescent cannabinoid exposure on neural and psychological processes. Neuroscience & Biobehavioral Reviews. 10.1016/j.neubiorev.2015.04.020

Iezzi, D., Cáceres-Rodríguez, A., Chavis, P., & Manzoni, O. J. (2025). Sex-specific disruptions in the developmental trajectory of anxiety-like behaviors due to prenatal cannabidiol exposure. Translational Psychiatry, 15(1), 354. 10.1038/s41398-025-03517-x

Iezzi, D., Caceres-Rodriguez, A., Chavis, P., & Manzoni, O. J. J. (2022). In utero exposure to cannabidiol disrupts select early-life behaviors in a sex-specific manner. Translational Psychiatry, 12(1), 501. 10.1038/s41398-022-02271-8

Innocenzi, E., De Domenico, E., Ciccarone, F., Zampieri, M., Rossi, G., Cicconi, R., Bernardini, R., Mattei, M., & Grimaldi, P. (2019). Paternal activation of CB2 cannabinoid receptor impairs placental and embryonic growth via an epigenetic mechanism. Scientific Reports, 9(1), 17034. 10.1038/s41598-019-53579-3

Jenkins, B. W., Moore, C. F., Jantzie, L. L., & Weerts, E. M. (2025). Prenatal cannabinoid exposure and the developing brain: Evidence of lasting consequences in preclinical rodent models. Neuroscience and Biobehavioral Reviews, 175, 106207. 10.1016/j.neubiorev.2025.106207

Katsidoni, V., Kastellakis, A., & Panagis, G. (2013). Biphasic effects of Δ9-tetrahydrocannabinol on brain stimulation reward and motor activity. International Journal of Neuropsychopharmacology, 16(10), 2273–2284. 10.1017/S1461145713000709

Koto, P., Allen, V. M., Fahey, J., & Kuhle, S. (2022). Maternal cannabis use during pregnancy and maternal and neonatal outcomes: A retrospective cohort study. BJOG: An International Journal of Obstetrics & Gynaecology, 129(10), 1687–1694. 10.1111/1471-0528.17114

Kuroda, K. O., & Tsuneoka, Y. (2013). Assessing Postpartum Maternal Care, Alloparental Behavior, and Infanticide in Mice: With Notes on Chemosensory Influences. En K. Touhara (Ed.), Pheromone Signaling (Vol. 1068, pp. 331–347). Humana Press. 10.1007/978-1-62703-619-1_25

Lauer, J., Zhou, M., Ye, S., Menegas, W., Schneider, S., Nath, T., Rahman, M. M., Di Santo, V., Soberanes, D., Feng, G., Murthy, V. N., Lauder, G., Dulac, C., Mathis, M. W., & Mathis, A. (2022). Multi-animal pose estimation, identification and tracking with DeepLabCut. Nature Methods, 19(4), 496–504. 10.1038/s41592-022-01443-0

Lisk, R. D., Pretlow, R. A., & Friedman, S. M. (1969). Hormonal stimulation necessary for elicitation of maternal nest-building in the mouse (Mus musculus). Animal Behaviour, 17(4), 730–737. 10.1016/s0003-3472(69)80020-5

Maciel, I. D. S., Abreu, G. H. D. D., Johnson, C. T., Bonday, R., Bradshaw, H. B., Mackie, K., & Lu, H.-C. (2022). Perinatal CBD or THC Exposure Results in Lasting Resistance to Fluoxetine in the Forced Swim Test: Reversal by Fatty Acid Amide Hydrolase Inhibition. Cannabis and Cannabinoid Research, 7(3), 318–327. 10.1089/can.2021.0015

Metz, T. D., Allshouse, A. A., Hogue, C. J., Goldenberg, R. L., Dudley, D. J., Varner, M. W., Conway, D. L., Saade, G. R., & Silver, R. M. (2017). Maternal marijuana use, adverse pregnancy outcomes, and neonatal morbidity. American Journal of Obstetrics and Gynecology, 217(4), 478.e1–478.e8. 10.1016/j.ajog.2017.05.050

Navarrete, F., García-Gutiérrez, M. S., Gasparyan, A., Austrich-Olivares, A., Femenía, T., & Manzanares, J. (2020). Cannabis Use in Pregnant and Breastfeeding Women: Behavioral and Neurobiological Consequences. Frontiers in Psychiatry, 11. 10.3389/fpsyt.2020.586447

Nowak, R., Porter, R. H., Lévy, F., Orgeur, P., & Schaal, B. (2000). Role of mother-young interactions in the survival of offspring in domestic mammals. Reviews of Reproduction, 5(3), 153–163. 10.1530/ror.0.0050153

Ochiai, W., Kitaoka, S., Kawamura, T., Hatogai, J., Harada, S., Iizuka, M., Ariumi, M., Takano, S., Nagai, T., Sasatsu, M., & Sugiyama, K. (2021). Maternal and Fetal Pharmacokinetic Analysis of Cannabidiol during Pregnancy in Mice. Drug Metabolism and Disposition, 49(4), 337–343. 10.1124/dmd.120.000270

Okabe, S., Nagasawa, M., Kihara, T., Kato, M., Harada, T., Koshida, N., Mogi, K., & Kikusui, T. (2013). Pup odor and ultrasonic vocalizations synergistically stimulate maternal attention in mice. Behavioral Neuroscience, 127(3), 432–438. 10.1037/a0032395

Oke, S. L., Lee, K., Papp, R., Laviolette, S. R., & Hardy, D. B. (2021). In Utero Exposure to Δ9-Tetrahydrocannabinol Leads to Postnatal Catch-Up Growth and Dysmetabolism in the Adult Rat Liver. International Journal of Molecular Sciences, 22(14), 7502. 10.3390/ijms22147502

Parolin, M., & Simonelli, A. (2016). Attachment Theory and Maternal Drug Addiction: The Contribution to Parenting Interventions. Frontiers in Psychiatry, 7, 152. 10.3389/fpsyt.2016.00152

Parra-Vargas, M., Ramon-Krauel, M., Lerin, C., & Jimenez-Chillaron, J. C. (2020). Size Does Matter: Litter Size Strongly Determines Adult Metabolism in Rodents. Cell Metabolism, 32(3), 334–340. 10.1016/j.cmet.2020.07.014

Rokeby, A. C. E., Natale, B. V., & Natale, D. R. C. (2023). Cannabinoids and the placenta: Receptors, signaling and outcomes. Placenta, 135, 51–61. 10.1016/j.placenta.2023.03.002

Sarrafpour, S., Urits, I., Powell, J., Nguyen, D., Callan, J., Orhurhu, V., Simopoulos, T., Viswanath, O., Kaye, A. D., Kaye, R. J., Cornett, E. M., & Yazdi, C. (2020). Considerations and Implications of Cannabidiol Use During Pregnancy. Current Pain and Headache Reports, 24(7), 38. 10.1007/s11916-020-00872-w

Scheyer, A., Melis, M., Trezza, V., & Manzoni, O. J. J. (2019). Consequences of Perinatal Cannabis Exposure. Trends in Neurosciences, 42(12), 871–884. 10.1016/j.tins.2019.08.010

Schmid, P. C., Paria, B. C., Krebsbach, R. J., Schmid, H. H. O., & Dey, S. K. (1997). Changes in anandamide levels in mouse uterus are associated with uterine receptivity for embryo implantation. Proceedings of the National Academy of Sciences of the United States of America, 94(8), 4188–4192. 10.1073/pnas.94.8.4188

Schuster, L., Henderson, R., Sankar, S., Ananth, D., Kirk, M., Leone, P., Adolph, K. E., Froemke, R. C., & Mar, A. (2022). Longitudinal monitoring of maternal care and maternal neglect. Preprint. 10.1101/2022.12.26.521927

Smart, J. L. (1976). Maternal behaviour of undernourished mother rats towards well fed and underfed young. Physiology & Behavior, 16(2), 147–149. 10.1016/0031-9384(76)90298-5

Swenson, K. S., Gomez Wulschner, L. E., Hoelscher, V. M., Folts, L., Korth, K. M., Oh, W. C., & Bates, E. A. (2023). Fetal cannabidiol (CBD) exposure alters thermal pain sensitivity, problem-solving, and prefrontal cortex excitability. Molecular Psychiatry, 28(8), 3397–3413. 10.1038/s41380-023-02130-y

Tagawa, N., Tsuneoka, Y., & Funato, H. (2025). Nest-building behavior in laboratory mice: Multifunctional roles and neural mechanisms. Neuroscience & Biobehavioral Reviews, 178, 106377. 10.1016/j.neubiorev.2025.106377

Topilko, T., Diaz, S. L., Pacheco, C. M., Verny, F., Rousseau, C. V., Kirst, C., Deleuze, C., Gaspar, P., & Renier, N. (2022). Edinger-Westphal peptidergic neurons enable maternal preparatory nesting. Neuron, 110(8), 1385–1399.e8. 10.1016/j.neuron.2022.01.012

Traccis, F., Frau, R., & Melis, M. (2020). Gender Differences in the Outcome of Offspring Prenatally Exposed to Drugs of Abuse. Frontiers in Behavioral Neuroscience, 14, 72. 10.3389/fnbeh.2020.00072

Vanin, S. R., Lee, K., Nashed, M., Tse, B., Sarikahya, M., Brar, S., Tomy, G., Lucas, A.-M., Tomy, T., Laviolette, S. R., Arany, E. J., & Hardy, D. B. (2023). Gestational exposure to cannabidiol leads to glucose intolerance in 3-month-old male offspring. Journal of Endocrinology, 260(1), e230173. 10.1530/JOE-23-0173

Volkow, N. D., Han, B., Compton, W. M., & McCance-Katz, E. F. (2019). Self-reported Medical and Nonmedical Cannabis Use Among Pregnant Women in the United States. JAMA, 322(2), 167–169. 10.1001/jama.2019.7982

Wang, H., Guo, Y., Wang, D., Kingsley, P. J., Marnett, L. J., Das, S. K., DuBois, R. N., & Dey, S. K. (2004). Aberrant cannabinoid signaling impairs oviductal transport of embryos. Nature Medicine, 10(10), 1074–1080. 10.1038/nm1104

Weber, E. M., & Olsson, I. A. S. (2008). Maternal behaviour in Mus musculus sp.: An ethological review. Applied Animal Behaviour Science, 114(1-2), 1–22. 10.1016/j.applanim.2008.06.006

Winters, C., Gorssen, W., Ossorio-Salazar, V. A., Nilsson, S., Golden, S., & D’Hooge, R. (2022). Automated procedure to assess pup retrieval in laboratory mice. Scientific Reports, 12(1), 1663. 10.1038/s41598-022-05641-w

Winters, C., Gorssen, W., Wöhr, M., & D’Hooge, R. (2023). BAMBI: A new method for automated assessment of bidirectional early-life interaction between maternal behavior and pup vocalization in mouse dam-pup dyads. Frontiers in Behavioral Neuroscience, 17, 1139254. 10.3389/fnbeh.2023.1139254

Wolterink-Donselaar, I. G., Meerding, J. M., & Fernandes, C. (2009). A method for gender determination in newborn dark pigmented mice. Lab Animal, 38(1), 35–38. 10.1038/laban0109-35

Wu, Y., Kirillov, A., Massa, F., Lo, W.-Y., & Girshick, R. (2019). *Detectron2* [Software]. https://github.com/facebookresearch/detectron2

Young-Wolff, K. C., Sarovar, V., Tucker, L.-Y., Avalos, L. A., Alexeeff, S., Conway, A., Armstrong, M. A., Weisner, C., Campbell, C. I., & Goler, N. (2019). Trends in marijuana use among pregnant women with and without nausea and vomiting in pregnancy, 2009 to 2016*. Drug and alcohol dependence, 196, 66–70. 10.1016/j.drugalcdep.2018.12.009

Zhang, Z., Dawson, P. A., Piper, M., & Simmons, D. G. (2019). Postnatal N-acetylcysteine administration rescues impaired social behaviors and neurogenesis in Slc13a4 haploinsufficient mice. EBioMedicine, 43, 435–446. 10.1016/j.ebiom.2019.03.081

Zimmer, M. R., Fonseca, A. H. O., Iyilikci, O., Pra, R. D., & Dietrich, M. O. (2019). Functional Ontogeny of Hypothalamic Agrp Neurons in Neonatal Mouse Behaviors. Cell, 178(1), 44–59.e7. 10.1016/j.cell.2019.04.026

