## Supplementary figures tables and legends for "Automated pup-level analysis reveals distinct effects of prenatal CBD and THC exposure on maternal retrieval"

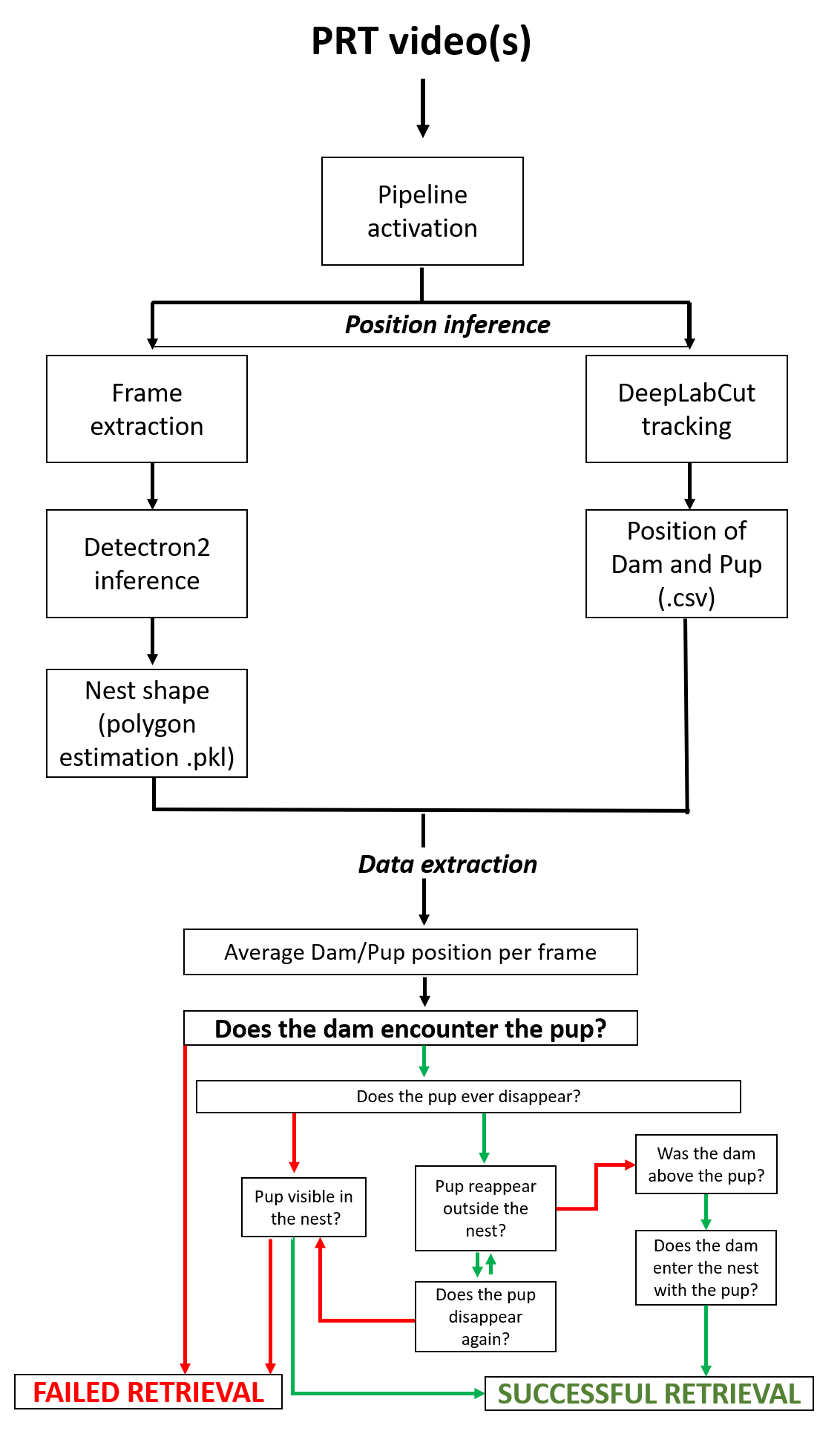

**Supplementary Figure 1: Schema of the pipeline required for the automated analysis.** Once the video is uploaded in the software, two different position extractions are performed: nest area detection and two-animal tracking. The positional data is used to determine whether the dam has encountered the pup and the frame in which it has been retrieved.

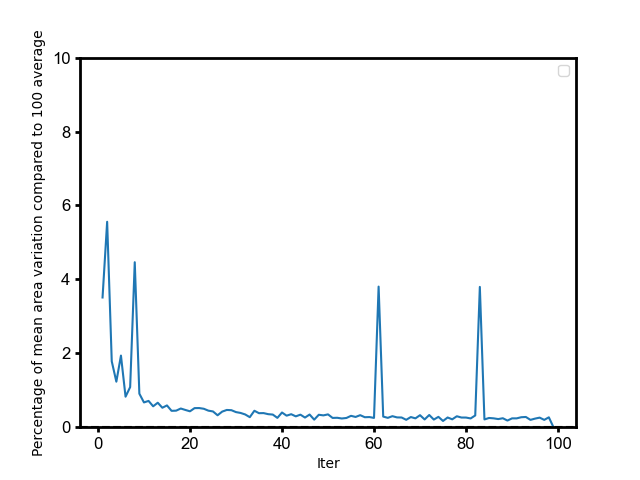

**Supplementary Figure 2: Effect on accuracy detection of number of frames used to estimate nest area**. This analysis evaluates how the number of video frames used to draw the nest mask influences the accuracy of nest area estimation. A reference value was established by averaging measurements across 100 frames. The graph plots the percentage difference between estimates obtained with fewer frames and this reference value. The X-axis shows the number of frames included in the calculation, while the Y-axis shows the deviation (%) from the reference area. As the number of frames increases, the estimates converge toward the reference, indicating that larger frame samples improve stability and accuracy; however, **20 frames were sufficient for correct estimation.**

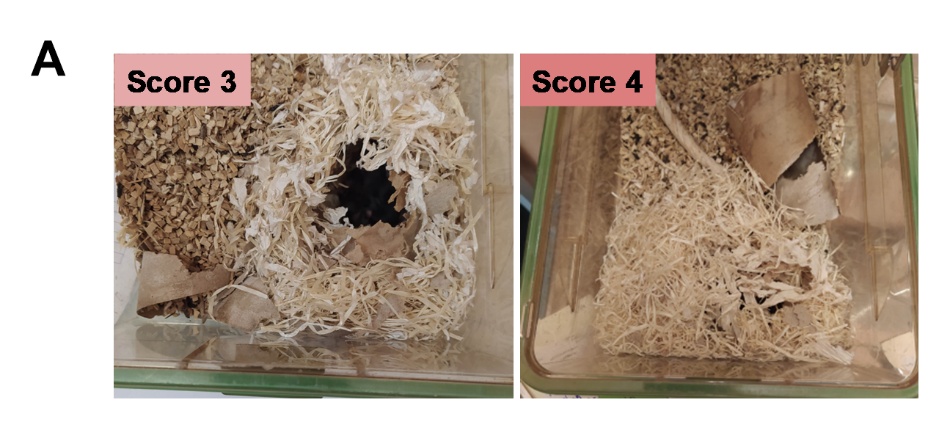

**Supplementary Figure 3: Home-cage nest grading** (A) Representative images illustrate the two most frequent nest quality scores: Score 3 corresponds to fully shredded nesting material forming walls tall enough to cover the dam; Score 4 reflects a complete enclosure with a roof, fully covering the dam.

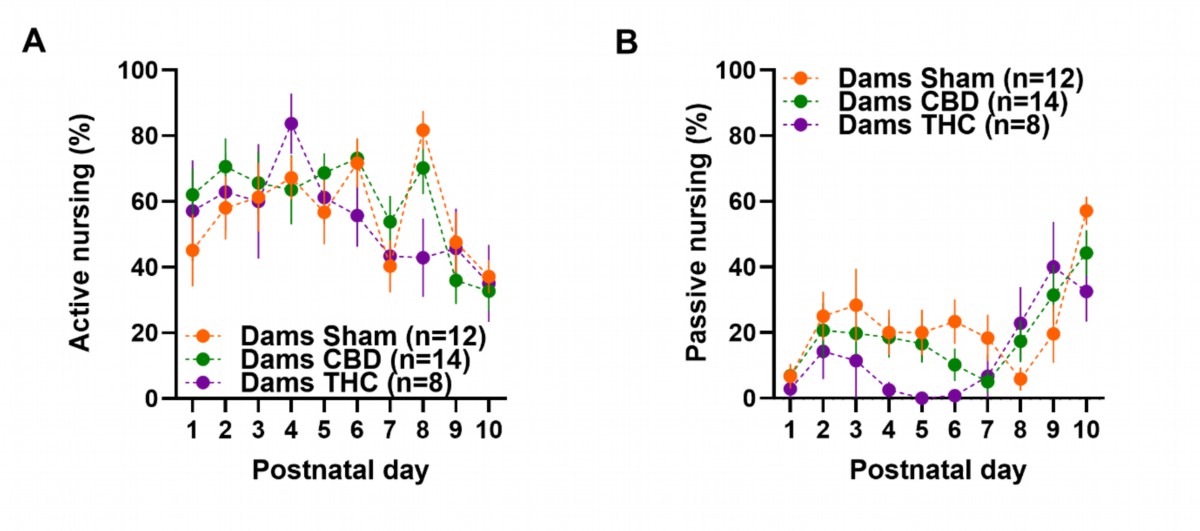

**Supplementary Figure 4: Nursing patterns during the first 10 postnatal days.** Nursing style shifts over time. Specifically, **(A)** active nursing predominates in the early postpartum period and transitions to **(B)** passive nursing by the end of the follow-up (**PND 9–10**). **(A–B)** Time-course data are presented as mean ± SEM in XY plots. Group sizes: Sham (n = 12), CBD (n = 14), THC (n = 8). Statistical analysis was performed via **mixed-effects models** with the Geisser-Greenhouse correction, followed by Tukey’s post hoc multiple comparison test. Only results with p < 0.05 were considered statistically significant and are reported.

| **Treatment** | **Levene Test** | **Trial #** | **N** | **Mean Weight (g)** | **Std. Deviation** | **MIN** | **MAX** |
| --- | --- | --- | --- | --- | --- | --- | --- |
| **SHAM** | 0.981 | 1 | 13 | 2.92 | 0.4 | 2.4 | 3.6 |
|  |  | 2 | 13 | 2.92 | 0.38 | 2.2 | 3.6 |
|  |  | 3 | 13 | 2.96 | 0.39 | 2.1 | 3.5 |
|  |  | 4 | 13 | 2.89 | 0.38 | 2.2 | 3.5 |
|  |  | 5 | 12 | 2.98 | 0.34 | 2.5 | 3.6 |
|  |  | 6 | 10 | 3.03 | 0.33 | 2.6 | 3.6 |
|  |  | 7 | 7 | 2.99 | 0.42 | 2.5 | 3.8 |
|  |  | 8 | 1 | 3.6 | — | 3.6 | 3.6 |
| **CBD** | 0.596 | 1 | 13 | 2.46 | 0.3 | 1.8 | 3 |
|  |  | 2 | 13 | 2.52 | 0.38 | 2 | 3.3 |
|  |  | 3 | 13 | 2.52 | 0.24 | 2.1 | 2.9 |
|  |  | 4 | 13 | 2.58 | 0.24 | 2.1 | 3 |
|  |  | 5 | 13 | 2.47 | 0.22 | 2.1 | 2.8 |
|  |  | 6 | 12 | 2.58 | 0.25 | 2.1 | 3 |
|  |  | 7 | 9 | 2.53 | 0.23 | 2.2 | 2.8 |
|  |  | 8 | 7 | 2.57 | 0.14 | 2.4 | 2.8 |
|  |  | 9 | 5 | 2.6 | 0.29 | 2.4 | 3.1 |
|  |  | 10 | 2 | 2.4 | 0.42 | 2.1 | 2.7 |
| **THC** | 0.994 | 1 | 8 | 2.44 | 0.36 | 2 | 3 |
|  |  | 2 | 8 | 2.46 | 0.39 | 2 | 3.2 |
|  |  | 3 | 8 | 2.41 | 0.38 | 1.8 | 3.1 |
|  |  | 4 | 8 | 2.41 | 0.36 | 1.8 | 2.9 |
|  |  | 5 | 8 | 2.34 | 0.35 | 1.9 | 2.9 |
|  |  | 6 | 6 | 2.5 | 0.34 | 2.1 | 3 |
|  |  | 7 | 6 | 2.63 | 0.33 | 2.2 | 3.1 |
|  |  | 8 | 6 | 2.38 | 0.29 | 1.9 | 2.7 |
|  |  | 9 | 4 | 2.33 | 0.33 | 2.1 | 2.8 |
|  |  | 10 | 2 | 2.3 | 0.14 | 2.2 | 2.4 |

**Supplementary Table 1:** **The random selection of pup order for the PRT task is not biased by body weight.** Levene's test indicated no significant differences in weight variability based on the order in which pups from the same litter were tested. The table summarizes pup weight statistics for each treatment group, organized by testing order. Reported values include treatment group, Levene's test p-value, testing order, number of pups (n), mean pup weight (g), standard deviation (std deviation), and minimum (MIN) and maximum (MAX) pup weights.
